# Nucleotide-free structures of Kif20A illuminate the atypical allostery in this mitotic kinesin-6

**DOI:** 10.1101/2022.11.10.515925

**Authors:** Fanomezana Moutse Ranaivoson, Vincent Crozet, Matthieu P.M.H. Benoît, Amna Abdalla Mohammed Khalid, Carlos Kikuti, Helena Sirkia, Ahmed El Marjou, Ana B. Asenjo, Hernando Sosa, Christoph F. Schmidt, Steven S. Rosenfeld, Anne Houdusse

## Abstract

KIF20A is a critical kinesin for cell division and a promising anti-cancer drug target. The mechanisms underlying its cellular roles remain elusive. Interestingly, unusual coupling between the nucleotide- and microtubule-binding sites of this kinesin-6 has been reported but little is known about how its divergent sequence leads to atypical motility properties. We present here the first high-resolution structure of its motor domain that delineates the highly unusual structural features of this motor, including a long L6 insertion that integrates into the core of the motor domain and that drastically affects allostery and ATPase activity. Together with the high-resolution cryo-EM microtubule-bound KIF20A structure that reveal the microtubule-binding interface, we dissect the peculiarities of the KIF20A sequence that work to favor fast dissociation of ADP, particularly in contrast to other kinesins. Structural and functional insights from the KIF20A pre-power stroke conformation thus highlight the role of extended insertions in shaping the motor mechanochemical cycle. Essential for force production and processivity is the length of the neck linker in kinesins. We highlight here the role of the sequence preceding the neck linker in controlling its backward docking and show that a neck linker 4-times longer than kinesin-1 is required for the activity of this motor.

## Introduction

Cell division requires the collective action of motor proteins in a precisely regulated manner, including several kinesins (Kin) that drive key steps in mitosis (Kozielski, 2015). Among these, the Kin-6 KIF20A, (MKLP2, Rabkinesin-6) plays critical roles during the metaphase/anaphase transition and cytokinesis (Fontijn *et al*., 2001; Hill *et al*., 2000). KIF20A is a plus end directed motor that was initially identified as a RAB6 effector that promotes dissociation of RAB6-positive vesicles from Golgi membranes by partnering with myosin-2 (Echard *et al*., 1998, 2001; Miserey-Lenkei *et al*., 2017). In mitosis, KIF20A triggers the translocation of the Chromosomal Passenger Complex (CPC) from the centromeres to the spindle midzone and its subsequent enrichment at the equatorial cortex to control furrow ingression and abscission (Gruneberg *et al*., 2004; Landino *et al*., 2017).

KIF20A is upregulated in a variety of malignancies, and its depletion reduces tumor cell viability (Gasnereau *et al*., 2012; Saito *et al*., 2017; Taniuchi *et al*., 2005; Yan *et al*., 2012; Z. Zhang *et al*., 2019), making it a target and biomarker for cancer therapies. KIF20A is a driver of castration-resistant prostate cancer progression (CRPC) via its role in vesicle secretion and is a promising therapeutic target against CRPC (Copello & Burnstein, 2022). Paprotrain, the first KIF20A pharmacologic inhibitor discovered (Tcherniuk *et al*., 2010), induces cytokinetic defects that lead to multi-nucleation. A potent derivative, BKS0349, induces apoptosis and inhibits cell proliferation in a xenograft murine model of endometriosis (Ferrero *et al*., 2019).

KIF20A is regulated by several effectors of the cell cycle including the master mitotic kinases Cdk1/Cyclin-B kinase (Hümmer & Mayer, 2009; Kitagawa *et al*., 2014), Aurora kinase B (Fung *et al*., 2017) and Polo-kinase 1 (Plk1) (Neef *et al*., 2003). Each of the three Kin-6 family members (KIF20A, KIF20B and KIF23) plays distinct roles in cytokinesis (Kozielski, 2015). KIF20A and KIF23 guide key mitotic regulators to their proper localization on the spindle midzone and in the midbody, *i.e.* the CPC (for KIF20A) (Gruneberg *et al*., 2004; Landino *et al*., 2017) or RacGAP1 (for KIF23) (Mishima *et al*., 2002; Pavicic-Kaltenbrunner *et al*., 2007), while the role of KIF20B is less clear (Janisch *et al*., 2018).

Despite the importance of KIF20A in mitosis and oncogenesis, little is known about its motor properties and its structure, although this information is critical to define the cellular role of the motor. Kif20A is reported to show minimal transport of low-load or high-load cargo in cells, unlike other mammalian Kin-6 members (Poulos *et al*., 2022). How the intriguing, atypical features of its motor domain sequence result in differences in the motor mechanism compared to other previously studied kinesins is currently unknown. KIF20A exhibits unusual kinetic signatures that distinguish it from other motors of the family including a relative uncoupling of the allosteric communication between the nucleotide binding site (NBS) and microtubule interface (Atherton *et al*., 2017). Among the kinesin classes, Kin-6 motor domains are unique in having long N- and C-terminal extensions as well as a particular long sequence inserted within loop L6. KIF20A also possesses a neck-linker (NL) four times longer than other kinesins (residues 506 to 553).

Previous studies in Kin-1 have shown that elongation of the NL reduces motor processivity (Andreasson *et al*., 2015; Yildiz *et al*., 2008). Yet KIF20A, despite its elongated NL, has been reported to move processively (Adriaans *et al*., 2020). Current knowledge about how other kinesin classes allow gating between their heads via restricted positions of their linkers therefore cannot explain how KIF20A functions.

Here we present a crystal structure of the KIF20A motor domain (MD) at 2.7 Å resolution in the nucleotide-free (NF) state. This structure provides a detailed description of unique sequence differences and their consequences for the structure and function of this atypical kinesin. In addition, a microtubule-bound KIF20A structure was determined using cryo-electron microscopy (cryo-EM) with an overall resolution of 3.1 Å for the microtubule and 3.4 Å for KIF20A. These structural insights concur in providing details on the changes occurring upon binding to the track and provide a framework for understanding the atypical kinetic properties of KIF20A we are reporting here and that are critical for understanding its atypical motor properties. An important aspect of this work is our functional characterization of the limits of the MD: the length and the role of both the NL and the unusually long N-terminal extension are characterized here. We show that the NL is oriented in the backward direction in the NF state and provide a detailed description of how this occurs. Taken together, this structural and mechanistic study provides an essential framework for deciphering the roles that this unconventional kinesin plays in cells.

## Results

### High resolution structures of the nucleotide-free KIF20A motor domain

We obtained a 2.7 Å resolution structure of a murine KIF20A fragment (residues 55-510) comprising the MD and the first residues of the NL (Figure 1A and Figure 1–figure supplement 1). This provides the first atomic-resolution model for this atypical kinesin (Figure 1–figure supplement 2, Supplementary files 1 and 3). Overall, the MD structure displays a typical kinesin fold (Kull *et al*., 1996) (Figure 1B and Figure 1–figure supplement 1) with the MD in the open conformation, which is generally observed when kinesins are bound to microtubule or tubulin (Benoit *et al*., 2021; Cao *et al*., 2014). Several loop insertions unique to KIF20A emerge from the surface of the core MD (Figure 1A and C). The most striking are in loop L6 (Gly192-Asp295) (Figure 1D), which contains a very long insertion of 99 residues when compared to the loop L6 of the Kin-1 KIF5B (Figure 1A and Figure 1–figure supplement 1). Such a large insertion in L6 is a feature common to all Kin-6 family members.

**Figure 1.**
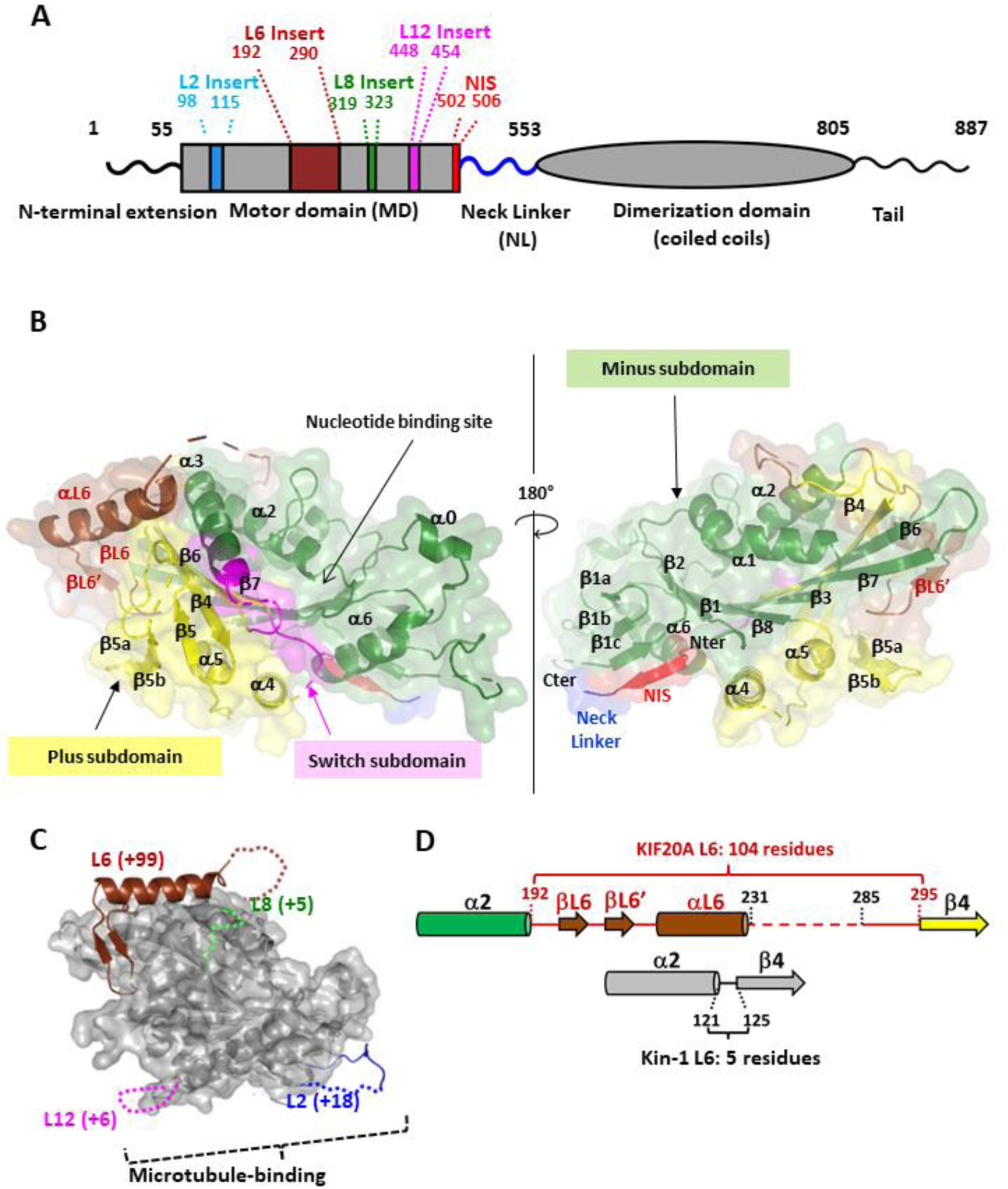
with 2 supplements. Structure of the KIF20A nucleotide-free motor domain. **(*A*)** Schematic of the mouse KIF20A highlighting important elements. **(*B*)** Crystal structure of the KIF20A MD with its three subdomains as defined in Benoit *et al*. 2021, shown in cartoon and semi-transparent surface mode, in two views. The structural elements that belong to the L6 insert (βL6, βL6’ and αL6) are shown in brown. The backward orientation of the “NL initial segment” (NIS) is highlighted (red). **(*C*)** Position and size of the KIF20A loop insertions. For each insertion, the number of KIF20A supplemental residues in comparison with the canonical Kin-1 MD fold is indicated in parentheses. **(*D*)** Diagram showing the structural organization of the 104 residue-long KIF20A L6 loop, compared to the 5 residue-long Kin-1 L6 loop.

To assess how microtubule binding influences this KIF20A NF conformation, we also determined the NF structure of the full-length MD, (1-565, Figure 2), while bound to microtubule by cryo-EM. This construct is extended at both the N- and C-terminus compared to the minimal 55-510 MD construct used in crystallography but it remains monomeric (Figure 2–figure supplement 1). 3D classification of the cryo-EM data resulted in two distinct classes (1 and 2) of KIF20A decorating microtubules, with one class (class-1) containing the majority of the particles (66%). Overall resolutions of 3.1 and 3.2 Å were obtained respectively for classes 1 and 2 (Figure 2–figure supplement 2), with 3.4 and 4.0 Å respectively for the kinesin part. The two classes differed mainly in the orientation of the MD relative to the microtubule so that the minus sub-domain in class-2 is positioned further away from the microtubule (Figure 2B). Also, the switch loops (L9 and L11) are well resolved in class-1 but not in class-2. Given the further separation from the microtubule interface and the lack of ordering of the switch loops, usually associated with microtubule binding in other kinesins (Atherton *et al*., 2017; Benoit *et al*., 2021; Shang *et al*., 2014), class-2 likely represents a partially unbound intermediate. In both classes the MD is in an open conformation.

**Figure 2.**
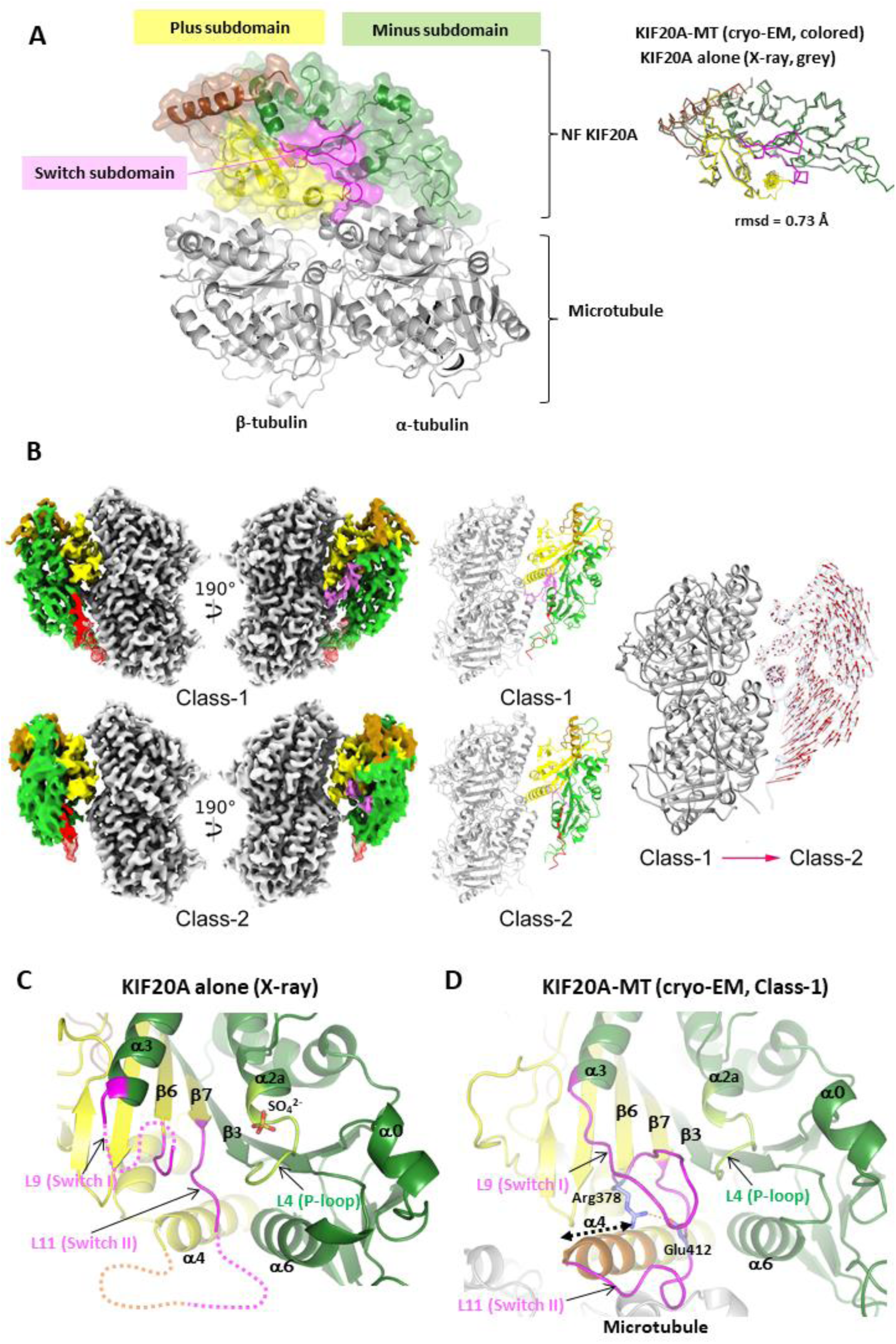
with 2 supplements. Structure of the KIF20A motor domain bound to microtubule and influence of microtubule to the nucleotide binding site. (*A*) Overall structure of NF KIF20A-microtubule complex, showing the three kinesin MD subdomains in cartoon and semi-transparent surface mode. The overall KIF20A conformation is similar to that of the MD alone determined by X-ray crystallography (rmsd = 0.73 Å, 288 aligned Cα atoms). (*B*) Cryo-EM density maps of the two classes of KIF20A-microtubule complexes rotated 190 degrees from each other (left panel) with corresponding fitted atomic models (center panel) and a comparison between the two class models (right panel). Regions of the maps and atomic models colored as in (A). In the right-most panel, the class-1 and class-2 structures are shown superimposed with the red vectors connecting equivalent Cα atoms from the class-1 to the class-2 structure. Note that relative to class-1, the class-2 KIF20A MD is rotated and the areas of the minus subdomain that make contacts with the microtubule in class-1 are further away from the microtubule surface in class-2. (*C*) The open NBS of KIF20A MD alone (X-ray structure) showing disordered switch loops. Close to the P-loop, a sulfate ion is found at the position normally occupied by the nucleotide β-phosphate. (*D*) The open NBS of KIF20A MD bound to microtubule, showing an ordering of the NBS loops and an extension of the α4 helix by three turns upon binding to microtubule. The conserved Arg378 and Glu412 are well positioned for a polar interaction (represented by a dotted line between ^Arg378^Nη1 and ^Glu412^Oε2).

The class-1 KIF20A-microtubule complex reveals a net ordering of several structural elements (Supplementary file 3), including microtubule-binding loops L2 and L12, which are longer in KIF20A than in other kinesins (Figure 1A and Figure 1–figure supplement 1 and Supplementary file 2), as well as the N-terminal end of helix α4. Interestingly, upstream to helix α4 the loop L11 (the switch II loop), which is partially disordered in the crystal structure, can be fully traced in the cryo-EM structure, and so is the loop L9 (the switch I loop) (Figure 2C and D). The N-terminally extended α4, together with the stabilized L11, serve as a platform for L9 to fold as a β-hairpin, bringing the conserved Arg378 (L9) and Glu412 (L11) to a position so that they can form a salt bridge (Figure 2C and D).

### The large specific L6 loop of KIF20A

The loop L6 (Gly192-Asp295) is composed of 104 residues in KIF20A, while in Kin-1 only 5 residues form this loop that links the α2b helix and the β4 strand (Figure 3A). Rather than extending entirely from the MD, our structures show that this fragment partly folds into it. The N-terminal and C-terminal regions (L6-N, 192-231; L6-C, 286-295) are structured and well defined (Figure 1–figure supplement 2). No density is visible for the central region of L6 (L6-M, Gln232-Pro285) both in crystallographic and cryo-EM density maps (Supplementary file 3) and its sequence also indicates that it contains flexible or intrinsically disordered regions. The L6-N forms a short β-hairpin and a α-helix (βL6, βL6’ and αL6) (Figure 1B, Figure 3A-C, Figure 1–figure supplement 1) that lies along a surface shared by the plus and minus subdomains (subdomains defined as in (Benoit *et al*., 2021)). The L6-C region (Asp286-Asp295) adopts an extended conformation between α1 and α2b (Figure 1–figure supplement 2). This L6-C region is less visible in the cryo-EM map, suggesting it is prone to higher flexibility.

**Figure 3.**
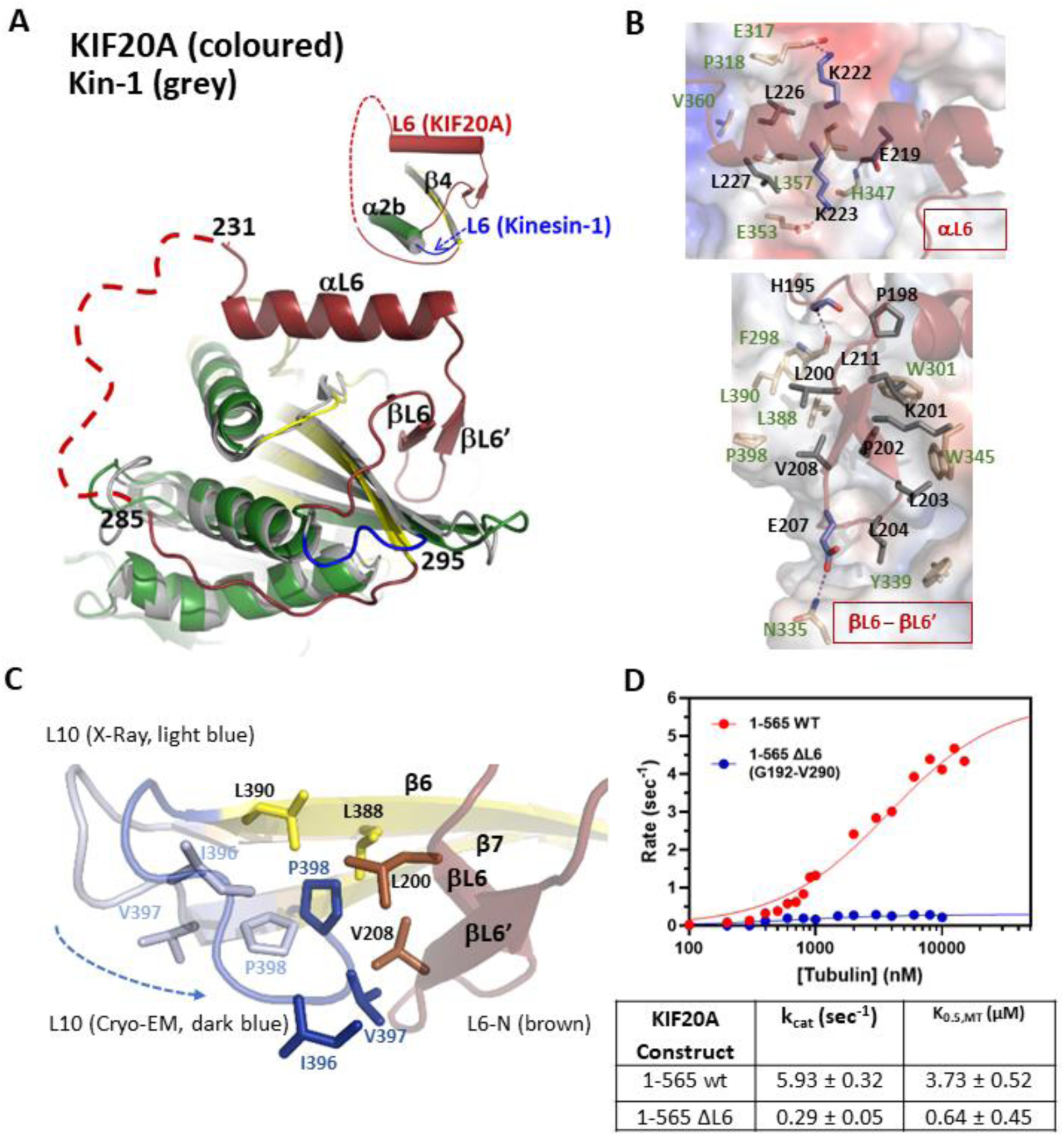
with 1 supplement. The KIF20A L6 insertion. **(*A*)** Superposition of KIF20A and Kin-1 (4LNU) MD structures highlighting the conformation of the 104-residue-long KIF20A L6 (brown), in comparison to the Kin-1 L6. **(*B*)** Interaction of the 192-232 (L6-N) fragment of the L6 insert with the core MD establishing several hydrophobic contacts with the core: the αL6 portion (E220, K222, K223) (left), and the βL6-βL6’ fragment (right). **(*C*)** Variation in the conformation of L10 between the X-ray and Cryo-EM structure, due to a crystal contact. I396, V397 and P398 are closer to V208 in the cryoEM structure. **(*D*)** microtubule-stimulated ATPase for monomeric 1-565 constructs, wild type (wt) or L6 insert-depleted (ΔL6: removal of 192-290 fragment).

The overall architecture and conformation of L6-N is similar in both X-ray and EM structure as it binds strongly to the MD core. The L6-N makes additional interactions with L10 in the cryo-EM map that are not possible in the crystal structure. The difference is due to a bias in the β6-L10-β7 conformation in the X-ray structure that is involved in crystal packing contacts (Figure 3–figure supplement 1A). The common interactions between L6-N and core MD residues are particularly well defined in the crystal structure, showing multiple hydrophobic contacts (Figure 3B), especially in the βL6-βL6’ region (Figure 3B, down). Hydrogen bonds and salt bridges (via residues ^αL6^Glu219, ^αL6^Lys222 and ^αL6^Lys223) are also involved in stabilizing interactions of the αL6 helix with the motor core (Figure 3B, up). A large surface, which is exposed in other kinesins, contains hydrophobic residues to accommodate L6-N in KIF20A (Figure 3B and arrows in Figure 1–figure supplement 1). In the cryo-EM structure, the closer proximity between the L10 loop and βL6-βL6’ hairpin appears to reinforce further these hydrophobic contacts, with the inclusion of ^L10^Ile396 and ^L10^Val397 in the ^L6^Val208 hydrophobic network (Figure 3C). Note that since most of these interactions are apolar, small variations in the L6-N side chain interactions with the MD core might be possible for adjusting potential rearrangements of the MD core during the motor cycle although they might influence the rate of these rearrangements. This long L6 element in Kin-6s thus provide constraints on the conformation of the central β-sheet that are quite distinct from those found in other kinesin subclasses, in which a 5-6 residues L6 linker connects the α2 helix and the β4 strand.

With minimal conformational adjustments such as those revealed by our comparison between the cryo-EM and the crystallographic structures presented here, the L6-N fragment likely follows the re-arrangements of the MD during the motor cycle. Furthermore, the L6-N beta hairpin region is sandwiched between the L8 loop at the interface with the microtubule on one side and the distal β6-L10-β7 region of the plus subdomain on the other side. These additional contacts made by L6-N with the β5a-β5b hairpin of the L8 loop (Figure 1–figure supplement 1 and Figure 3–figure supplement 2B) suggest that L6-N can influence the KIF20A association with microtubule or the rearrangements of the motor while bound to the track, since L8 is involved in microtubule binding (Atherton *et al*., 2017; Gigant *et al*., 2013). In addition, L6 provides a bridge between the plus and minus subdomains through a hydrophobic cluster between the L6-N residue Leu194 and residues from both subdomains (Figure 3–figure supplement 2C), indicating that L6 can influence the re-orientation of subdomains during motor function. Consistently, shortening of L6 to a 5-residue linker as in Kin-1 (delta L6, 192-290) drastically affects ATPase steady-state parameters of a fully active monomeric construct (1-565), with a 20-fold reduction of the k_cat._ (Figure 3D). As predicted from the structure, these kinetic studies confirm the role of L6 in the transient rearrangements of KIF20A during the motor cycle.

The long L6 loop is thus a structural element that plays an important role in the divergence of KIF20A mechano-chemistry from that of other kinesins. Our structures now provide the grounds for guiding detailed future investigations defining its precise role along the motor cycle and for the function of this kinesin in cells.

### KIF20A adopts a stable open conformation independent of microtubule binding

It is unusual for a kinesin to crystallize in the open state in absence of microtubule. Here we show that the overall KIF20A conformation is not affected by microtubules binding (rmsd = 0.73 Å of 286 aligned Cα between the crystal (MD alone) and the cryo-EM (microtubule-bound MD) structures) (Figure 2A) and that the NBS is open in both cases. Therefore microtubule-binding does not induce opening of the NBS as has been observed in other kinesins (Benoit et al., 2021; Hunter et al., 2022; Shang et al., 2014). Interestingly, Zen4, another Kin-6, has also been crystallized in a similar open conformation in the absence of tubulin or microtubule (Guan *et al*., 2017). We also found that addition of 5-10 mM Mg^2+^.ADP did not disrupt the ability to crystallize KIF20A in the same conformation. No nucleotide density or ordering of the NBS was visible in the electron density maps in this case. SAXS data however indicates that the scattering curve differs when 5 mM Mg^2+^.ADP is added (Figure 4–figure supplement 1). We also measured the affinity for Mg^2+^.ADP for this kinesin and found it to be ~5000 fold lower than for Kin-1 (Figure 4A, (Sadhu & Taylor, 1992)). Altogether, these data suggest that KIF20A likely does not stay in the open state in the presence of Mg^2+^.ADP when detached from microtubule, although the open state is particularly stabilized for this kinesin when no nucleotide is bound, even in the absence of microtubule.

**Figure 4.**
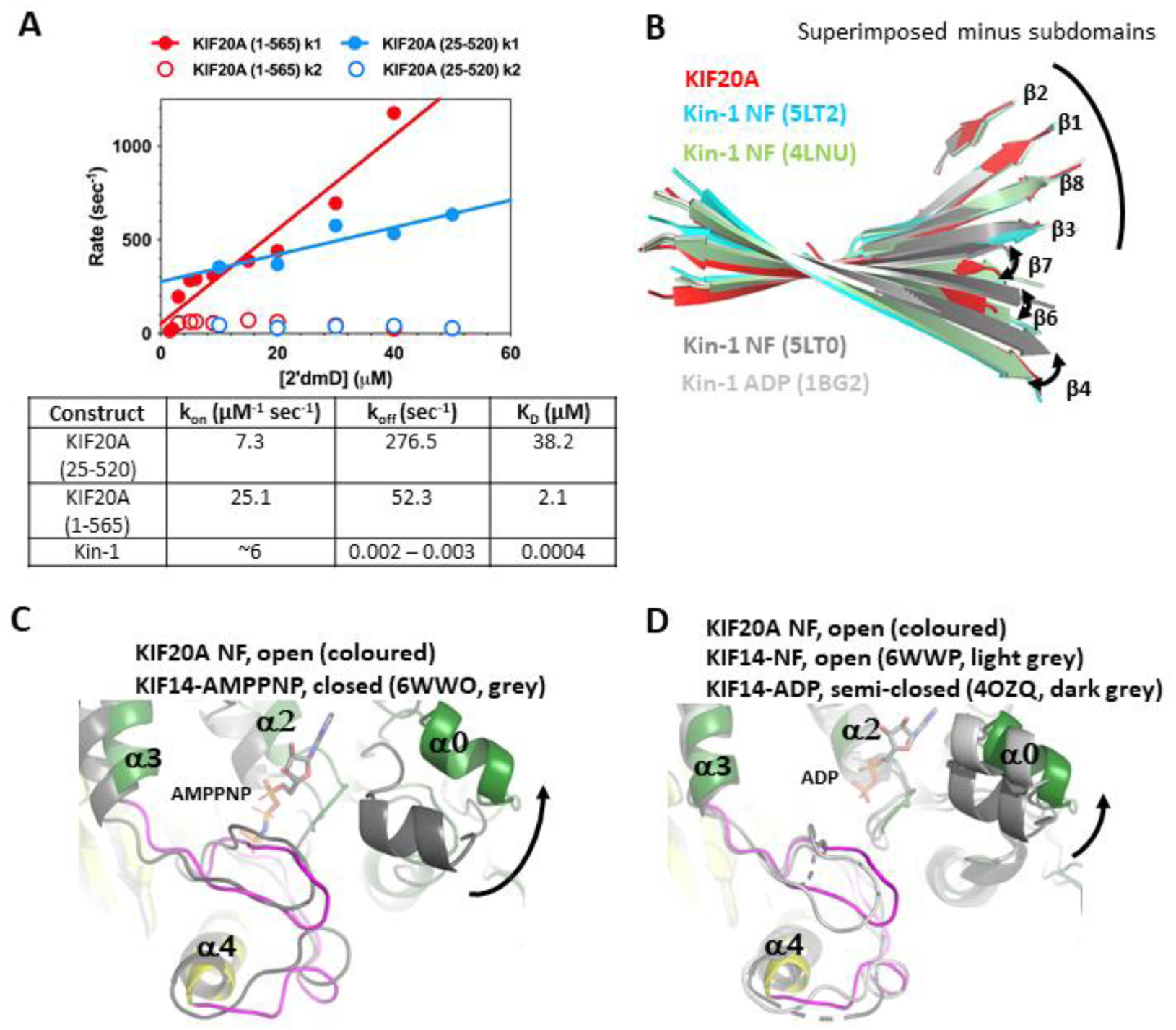
with 2 supplements. KIF20A exhibits a low affinity for ADP and an open nucleotide binding site (NBS). **(*A*)** Plots of rate constants for the fast (closed circles) and slow (open circles) phases of the biphasic transient produced by mixing 2’dmD with a 1-565 (red) and a 25-520 (blue) nucleotide free Kif20A MD construct. Linear fits of a plot of the 2’dmD concentration dependence of the fast phase provide measures of apparent second order rate constants for 2’dmD binding (**k_on_**), dissociation (**k_off_**), and apparent dissociation constant (**K_D_**). **(*B*)** Twist of the central β-sheet of KIF20A compared to that of other NF structures of Kin-1 (Cao *et al*., 2014, 2017). **(*C*, *D*)** Comparison of KIF20A with the monomeric closed state (C), semi-open, and open states (D) of the Kin-3 KIF14 (Arora *et al*., 2014; Benoit *et al*., 2021), showing a KIF20A NBS as open as in the microtubule-bound NF structure of KIF14. The microtubule-binding elements of the different MD structures were aligned for this comparison.

With our high-resolution structure, we next deciphered what could lead to this unusual stabilization of the open form, a conformation usually populated when kinesins are bound to microtubule. The highly exposed KIF20A NBS is associated with a β-sheet conformation among the most twisted in kinesins, a conformation similar to that of the tubulin-bound NF Kin-1 (4LNU, (Cao *et al*., 2014)) (Figure 4B), or to that of the microtubule-bound NF Kin-3 KIF14 (Benoit *et al*., 2021). Characteristic of these open conformations are a more twisted central β-sheet conformation and an outward position of the α0 helix (Figure 4C and D). Molecular dynamics simulations previously predicted that the α0 helix mobility influences ATP recruitment in (Hwang et al., 2017).

KIF20A-specific residues are likely to contribute to stabilize the open KIF20A NBS observed in our structures (Figure 4–figure supplement 2). For example, in Kin-1, the twist of the central β-sheet of the NF state is prone to explore at least two different conformations (Figure 4B, (Cao *et al*., 2017)), one of which corresponds to the conformation of the ADP-bound crystal structure (1BG2, (Kull *et al*., 1996)). In KIF20A, the β3 strand residue Trp153 appears to constrain the central β-sheet in a highly twisted conformation, most similar to the tubulin-bound Kin-1 NF structure (Figure 4–figure supplement 2I-H). A bulky residue at this position is only found in the KIF20A and KIF20B sequences while other kinesins have small side chain residues, which is more favourable for this transition. This could contribute to the preference of KIF20A for an open NBS state.

Likewise, the outward conformation of the α0 helix results from the specific sequence of KIF20A. In microtubule-free Kin-1 bound to ADP, the inward position of α0 is stabilized by several intramolecular interactions that cannot be established by the homolog residues in KIF20A (Figure 4–figure supplement 2E and F). In addition, the insertion of Glu82 in the loop L1 (Figure 1–figure supplement 1) is not compatible with a conformation of the upstream helix α0 similar to that of ADP-bound Kin-1, since that would bring charge hindrance or disruption of hydrophobic contacts and exposure of the corresponding residues (Figure 4–figure supplement 2F and G).

In addition, ^Kin-1^Arg190, which has been previously described to be part of the “Mg-network” (Chang *et al*., 2013), is important for an adequate stabilization the nucleotide-associated Mg^2+^ (Figure 4–figure supplement 2A). This residue corresponds to Gln365 in KIF20A, and its side chain is too short for stabilizing a strong Mg^2+^ coordinating network (Figure 4–figure supplement 2B). Therefore, Mg^2+^ and ADP is expected not to be as similarly attached to the KIF20A NBS as in Kin-1, which might explain the difference in affinity for Mg.ADP (Figure 4A). Likewise, ^Kin-1^Gln86 is involved in the conformational stabilisation of the P-loop (Figure 4–figure supplement 2C). Val160 occupies this position in KIF20A, which should result to a less firm anchoring of the P-loop by α6 residues and therefore a less stable association with the nucleotide (Figure 4–figure supplement 2D). In addition, the long L6 loop and the C-terminal extension could also participate in the stabilization of this open state.

In summary, as opposed to previously studied purely motile kinesins, KIF20A adopts a pre-stroke state in the absence of nucleotide in which the NBS is as open when bound to the microtubule as when detached from the microtubule. This property of KIF20A appear related to its very low ADP affinity when detached from the microtubule (Figure 4A). Our structures indicate that this specific property results from differences in key residues that influence the conformations of either the β-sheet or the NBS. Thus, unlike other kinesins, KIF20A does not need the assistance of the microtubule to trigger the opening of the NBS that is required for ADP release (Clancy *et al*., 2011; Moyer *et al*., 1998; Rosenfeld *et al*., 2005).

### KIF20A is a slow motor with striking specificities in the influence of microtubules for its allosteric conformational rearrangements

To get insights in how these peculiar features of KIF20A mechanistic elements translate into function, we performed surface-gliding assays at room temperature (23°C) with a His-tagged dimeric motor construct (1-665) (Figure 2–figure supplement 2), attached at high densities to a coverslip by anti-His antibodies (Figure 5A and Figure 5–figure supplement 1) and two controls were also examined: fast dimeric Kin-1 (*Neurospora crassa N*Kin-1) and Kin-3 (*Caenorhabditis elegans* UNC104 U653) constructs (Figure 5B-D, Figure 5–figure supplement 2, Videos 1-7). The KIF20A constructs produced gliding speeds (51 ± 5 nm/s) ~40 times slower than *N*Kin-1 or the Kin-3 UNC104 U653 constructs (2000 ± 100 nm/s and 1700 ± 100 nm/s, respectively, Figure 5B-D). Since ADP release is unusually fast in this motor (Figure 4A), this slow *in vitro* motor activity most likely results from the uncoupling of allosteric communication between nucleotide- and microtubule-binding events previously reported (Atherton *et al*., 2017), that our structures emphasize.

**Figure 5.**
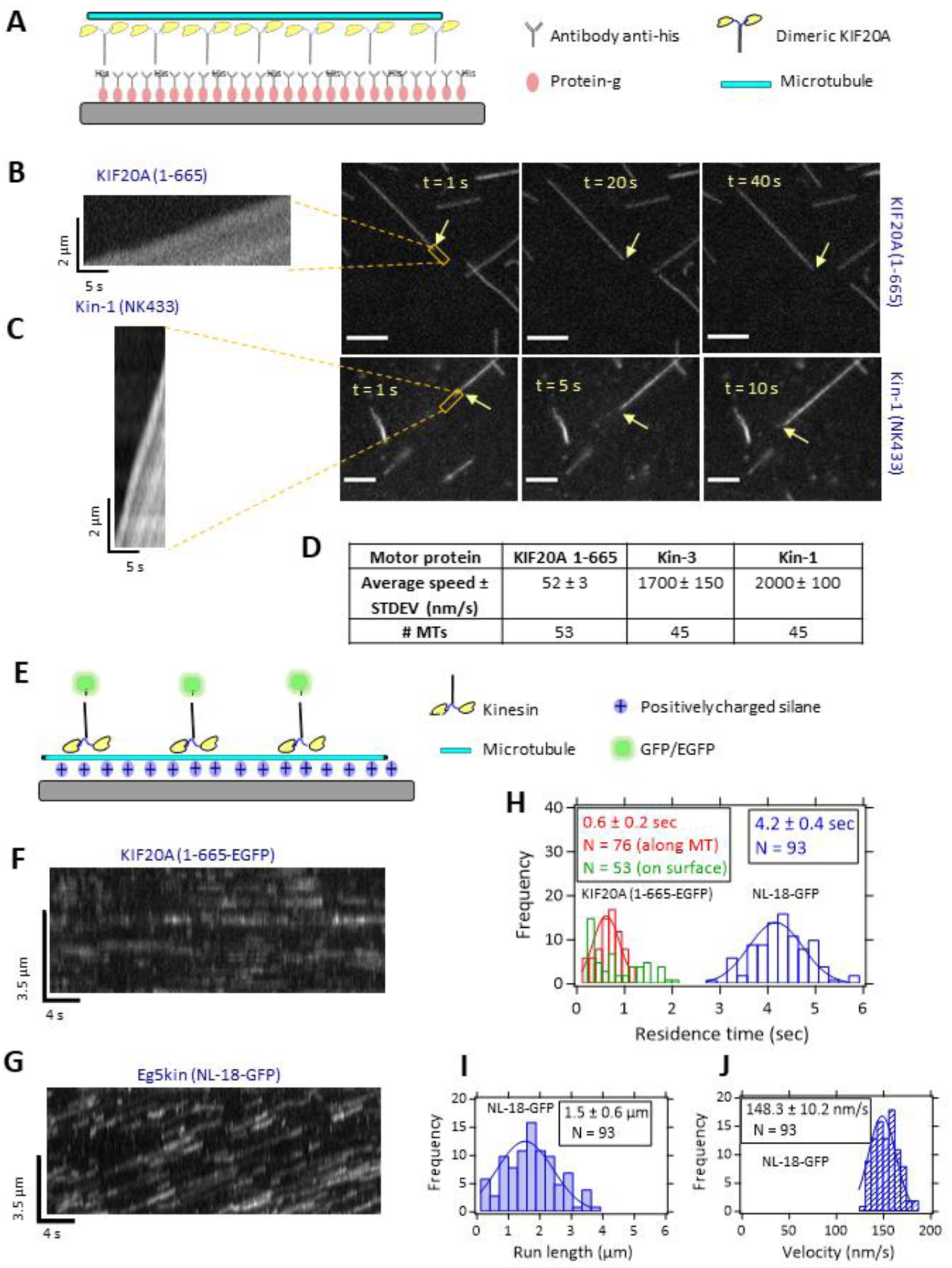
with 3 supplements. Surface gliding and single molecule motility of KIF20A 1-665. **(*A*)** Schematic: TMR-labeled microtubules were propelled by specifically attached motors. **(*B*)** Consecutive snapshots of TMR-labeled microtubules propelled by KIF20A 1-665 motors (Video 1) at saturating ATP (2 mM) (scale bars: 5 µm). A kymograph of single TMR-labeled microtubule propelled by collectives of KIF20A 1-665 is shown on the left. Yellow arrows point towards the end of the microtubule being tracked. **(*C*)** Consecutive snapshots of TMR-labeled microtubules propelled by Kin-1 (NK433) as control (scale bars: 5 µm). A kymograph of single TMR-labeled microtubule propelled by collectives of Kin-1 is shown on the left. All recordings at 10 frames per second. Orange rectangles show where kymographs were recorded. Microtubule-gliding velocities were determined by fitting kymographs (Figure 5–figure supplement 1A). Other constructs shown in Figure 5–figure supplement 2. **(*D*)** Average gliding speeds of KIF20A 1-665 and controls. **(E)** Schematic of the single-molecule TIRF experiment, in motility buffer, 2 mM ATP. Frame rate: 10 s^-1^. Kymographs generated in ImageJ. **(*F*)** Kymographs of 1-665-EGFP dimeric motors appearing on surface-immobilized microtubules (Video 11), bulk motor concentration: 20 nM (2 experiments). **(*G*)** GFP-labeled Kin-5 dimeric motors (Eg5kin-NL-18, see Methods) landing and moving on microtubules, bulk motor concentration 20 nM (2 experiments). **(*H*)** Single-molecule dwell times. **(*I, J*)** Run lengths and velocities, respectively, for Kin-5 control, measured from the kymograph in (*G*).

To determine if our tail-less dimeric KIF20A constructs are processive motors, we performed single-molecule fluorescence motility assays on surface-immobilized microtubules. The KIF20A (1-665-EGFP) construct moved microtubules in gliding assays (Figure 5–figure supplement 3 and Video 8). In single-molecule assays (Figure 5E), however, kymographs along a surface immobilized microtubules were indistinguishable from those recorded along a microtubule-free stretch of substrate next to a microtubule (Figure 5F, 5H and Video 9). We therefore could not detect binding or movement of individual motors. In the same assay, a tail-less dimeric Kin-5 motor used as control moved processively (Figure 5 G-J and Figure 5–figure supplement 3 and Video 10) as reported in the literature (Düselder *et al*., 2012). These results indicate that the dimeric KIF20A constructs missing the C-terminal tail are not processive. Note that it is common for non-processive motors to promote motility in gliding assays due to collective propulsion by many motors (Kaneko *et al*., 2020). These results are consistent with the data obtained with a human KIF20A tail-truncated construct (1-720) (Poulos *et al*., 2022). Only few events detected processivity using slow imaging rates and long imaging time.

We next compared the kinetics of ATP binding to 1-565 KIF20A MD using the fluorescent ATP analogue 2’ deoxy 3’ MANT-ATP (2’dmT). Data was acquired in the presence or absence of microtubules to probe whether the open NBS suggested by structural studies indeed facilitates binding of nucleotide compared to other kinesins. Mixing 2’dmT with NF preparations of MD or of a 1:4 MD-microtubule complex produced a biphasic fluorescence increase (Figure 6A for MD and 6B for MD-microtubule), defining fast and slow phases. The presence of two phases in the fluorescence transient implies two conformational changes occur following formation of a collision complex between the catalytic site and 2’dmT. Whether these steps occur sequentially or independently of each other is reflected in the dependence of the rate of each phase on ligand concentration (Rosenfeld & Taylor, 1984). For both MD (red) and MD-microtubule complexes (blue) (Figure 6C), the faster phase shows a clear hyperbolic dependency of rate on [2’dmT], defining maximum rates and apparent second order rate constants for 2’dmT binding, respectively, of 688.5 ± 92.0 s^-1^ and 169 µM^-1^.s^-1^ for MD; and 505 ± 42 s^-1^ and 193 µM^-1^.s^-1^ for MD-microtubule. The rate constant for the slower phase for MD shows no significant dependency on [2’dmT], as expected for sequential conformational changes, and averages 60-70 s^-1^ (Figure 6C, open red circles). While the rate of the slower phase for a MD-microtubule complex shows a very modest dependency on [2’dmT] at low ligand concentrations (Figure 6C, open blue circles), we conclude that these two conformational changes are likely sequential in a MD-microtubule complex as well.

**Figure 6.**
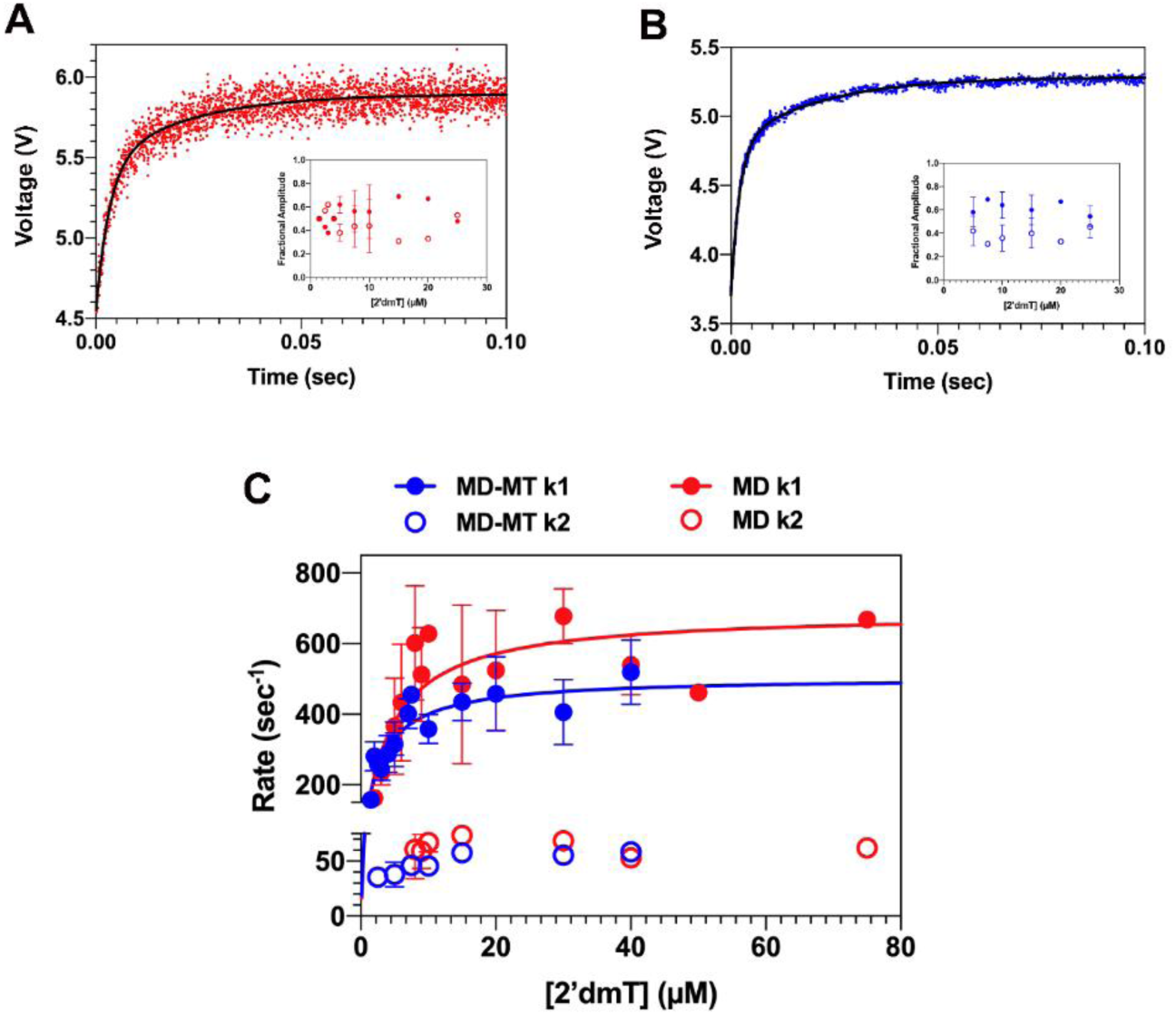
Kinetic properties of monomeric constructs of KIF20A, revealing the involvement of the N-terminal extension and long NL in its activity. (*A*) Fluorescence transient produced by mixing NF MD with 200 µM of 2’dmT. The fluorescence increases in two phases. The amplitudes of these two phases are depicted in the inset (closed red, fast phase; open red, slow phase), are nearly equal in magnitude and show little dependence on [2’dmT]. (*B*) Fluorescence transient produced by mixing {NF MD}:microtubule complex (1:4 concentration ratio) with 200 µM of the fluorescent ATP analogue 2’dmT. The fluorescence increases in two phases. The amplitudes of these two phases are depicted in the inset (closed blue, fast phase; open blue, slow phase), are nearly equal in magnitude and show little dependence on [2’dmT]. In (*A*) and (*B*), the relative amplitudes for the fast phase ranged from 40-60% of the total amplitude (closed circles in insets). (*C*) Plots of the binding rates of the fluorescent ATP analogue 2’dmT to the 1-565 KIF20A MD construct in the presence of microtubules (1:4 ratio (MD:microtubules), blue) and MD alone (red). Fitting to a bi-exponential rate equation shows rate constants k_1_ and k_2_ are similar and insensitive to the presence of microtubules (MD k_1_ = 688.5 ± 92.0 s^-1^; MD-microtubules k_1_ = 505 ± 42 s^-1^; MD k_2_ = 169 µM^-1^sec^-1^; MD-microtubules k_2_ = 193 µM^-1^s^-1^).

As we found for the shorter 25-520 construct (Atherton *et al*., 2017), microtubule binding has little effect on the kinetics of ATP binding to or release from the catalytic site of the 1-565 KIF20A MD. However, one important difference is that the apparent second-order rate constant for 2’dmT binding to the 1-565 MD is 10-20 fold faster than that for the shorter 25-520 MD version (Atherton *et al*., 2017), and 20-50 fold faster than corresponding values for Kin-1 or Eg5 (Cochran *et al*., 2004). This suggests that the catalytic site in NF KIF20A is considerably more accessible to nucleotide than in other kinesins. This is consistent with our finding that KIF20A adopts a stable NF state with a wide-open active site, as discussed above (Figure 4B). In addition, these results suggest a critical role of the N- and C-terminal extensions in KIF20A in stabilizing this NF conformation or in influencing the interaction with microtubule.

### Microtubule-binding interface – comparison with other kinesins

The KIF20A MD interacts with the microtubule through similar areas to those observed in other kinesin-microtubule complexes (L8, L11, H4, L12, H6). However, residue differences in these areas results in different conformations of the interacting regions, the contacts formed and the overall orientation of the MD relative to the microtubule (Figure 7). The areas of contact with tubulin (the “footprint”) are also similar to that observed in motile kinesins with most of the contacts made with the terminal helix-12 of α- and β-tubulin. However, KIF20A specific residues in these areas result in different contacts and a different orientation of the MD relative to tubulin (Figure 7C and D and Figure 7–figure supplement 1) as compared to other kinesins. This difference was mentioned by comparison of Kin-1 and low resolution KIF20A structures (Atherton *et al*., 2017). Here we show that the shift is even more pronounced when the microtubule-bound KIF20A is compared to Kin-3 (Figure 7B) (Benoit *et al*., 2021). In addition, to these interactions, the KIF20A elongated loop L2 and the minus-end pointing NL of this NF structure locate very near the microtubule interface and form salt bridges and hydrophobic interactions with α-tubulin (Figure 7C).

**Figure 7.**
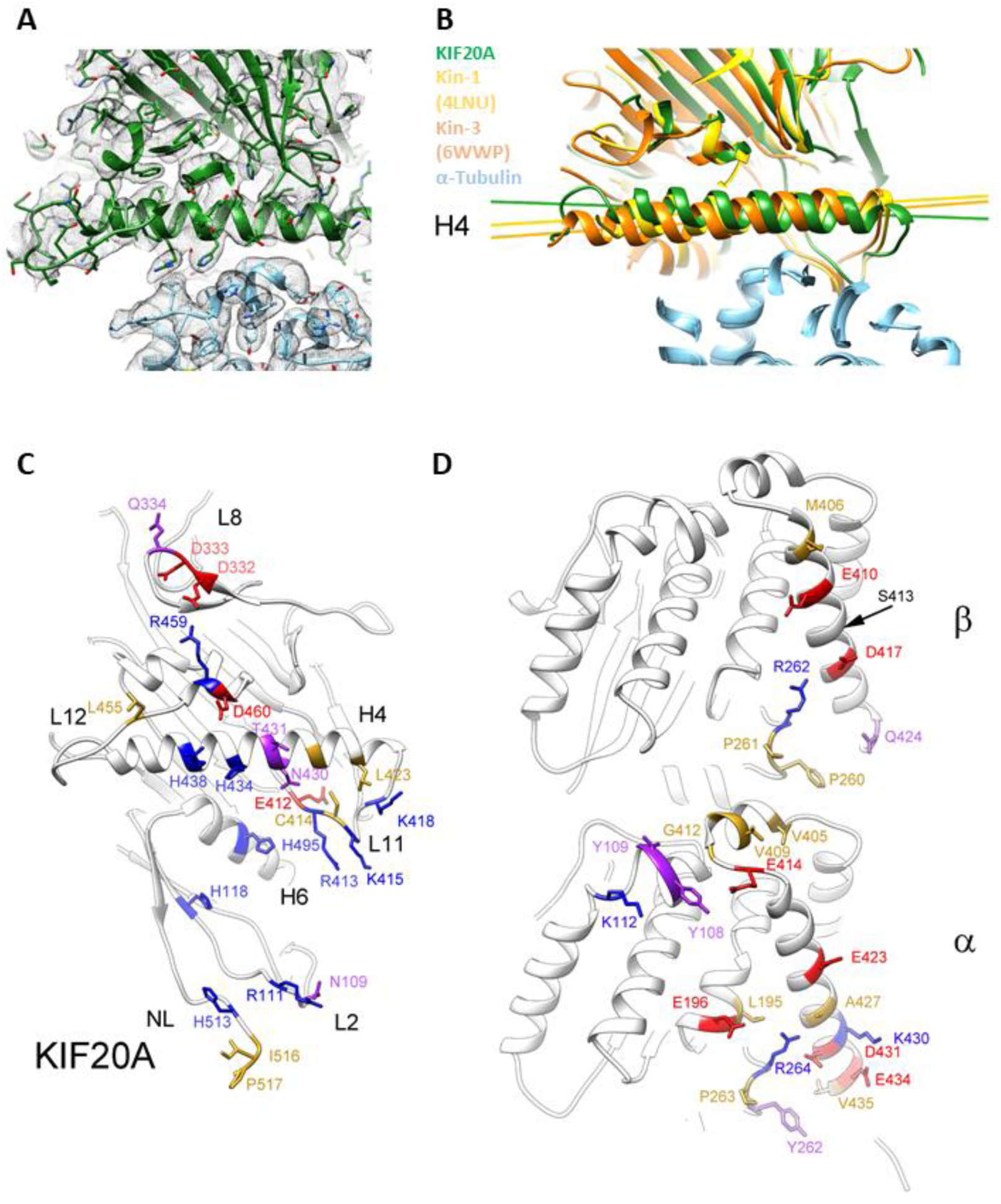
with 1 supplement. Interface Kinesin-Tubulin. **(*A*)** KIF20A-microtubule interface (class-1) near the KIF20A α-helix 4 (H4). KIF20A MD in green, tubulin in light blue and corresponding cryo-EM density isosurface as a semi-transparent grey mesh. **(*B*)** The KIF20A MD in the microtubule complex is rotated relative to the MDs in the Kin-1 and the Kin-3 microtubule complexes. All three microtubule or tubulin complex structures are aligned to their corresponding α-tubulin subunits. KIF20A MD in green, Kin-1 (KIF5B) MD in yellow, Kin-3 (KIF14) MD in orange, and α-tubulins in light blue. Views in (*A*) and (*B*) are from the microtubule plus end towards the minus end. **(*C*)** View of the KIF20A-microtubule interface from the microtubule. **(*D*)** View of the KIF20A-microtubule interface from the KIF20A MD. In (*C*) and (D) KIF20A and microtubule residues making contacts at the interface are shown as stick atom representations and colored according to the residue properties (negative in red, positive in blue, polar in purple and hydrophobic in yellow) Residues making contacts at the interface were identified using the UCSF-Chimera routine “find clashes/contacts” (Pettersen *et al*., 2004).

### The role of the NIS in tuning NL directionality

In both the X-ray and cryo-EM open structures, the KIF20A NL is oriented towards the microtubule minus-end, similar to what was previously described for Zen4 (Guan *et al*., 2017). In these Kin-6s, the “NL Initial Segment” (NIS) (Nitta *et al*., 2008) is docked onto the core MD by forming a β-strand interacting with β1c (Figures 1B, 8A, 8B, and Figure 8–figure supplement 1A). Interestingly, the structure of the ADP-bound Kin-10 KIF22 (6NJE, Walker *et al*., unpublished) also exhibits a similar NIS conformation, demonstrating that this backward NIS docking is not restricted to Kin-6s and can occur when ADP is bound in a semi-closed NBS state. For other kinesins, backward docking was in fact observed for the lead head only in the particular condition that two heads would be bound to microtubule. We thus analyzed what promoted backward docking in KIF20A independent of load.

The backward docking of the NIS leads to a C-terminal shortening of the α6 helix by ~1.5 turns as compared to post-power-stroke structures (Figure 8F). The unwound residues are from the NIS (L502-L506) (Nitta *et al*., 2008) and from a short preceding pre-NIS linker (F499-L502). The pre-NIS linker sequence is adjusted to allow docking of the NIS immediately following the α6 helix promoting the NIS interaction as a supplemental β-strand to the peripheral β-sheet made of β1a, β1b and β1c via five intra-strand hydrogen bonds (Figure 8B). Interestingly, the class-1 cryo-EM map indicates how the NL after _NIS_Leu506 continues to form apolar interactions with the MD for at least 9 residues (until Leu514). Gly515 and Pro517 promote a turn in the NL so that the following residues mainly form apolar contacts with the microtubule (Figure 7C). In most of the other known pre-power stroke plus-end kinesin MD structures (NF or ADP-bound with either an open or semi-closed NBS), this region stays helical until the end of the NIS (^KIF5B^Lys323 which would correspond to ^KIF20A^Ser504) (Figure 8–figure supplement 1A).

**Figure 8.**
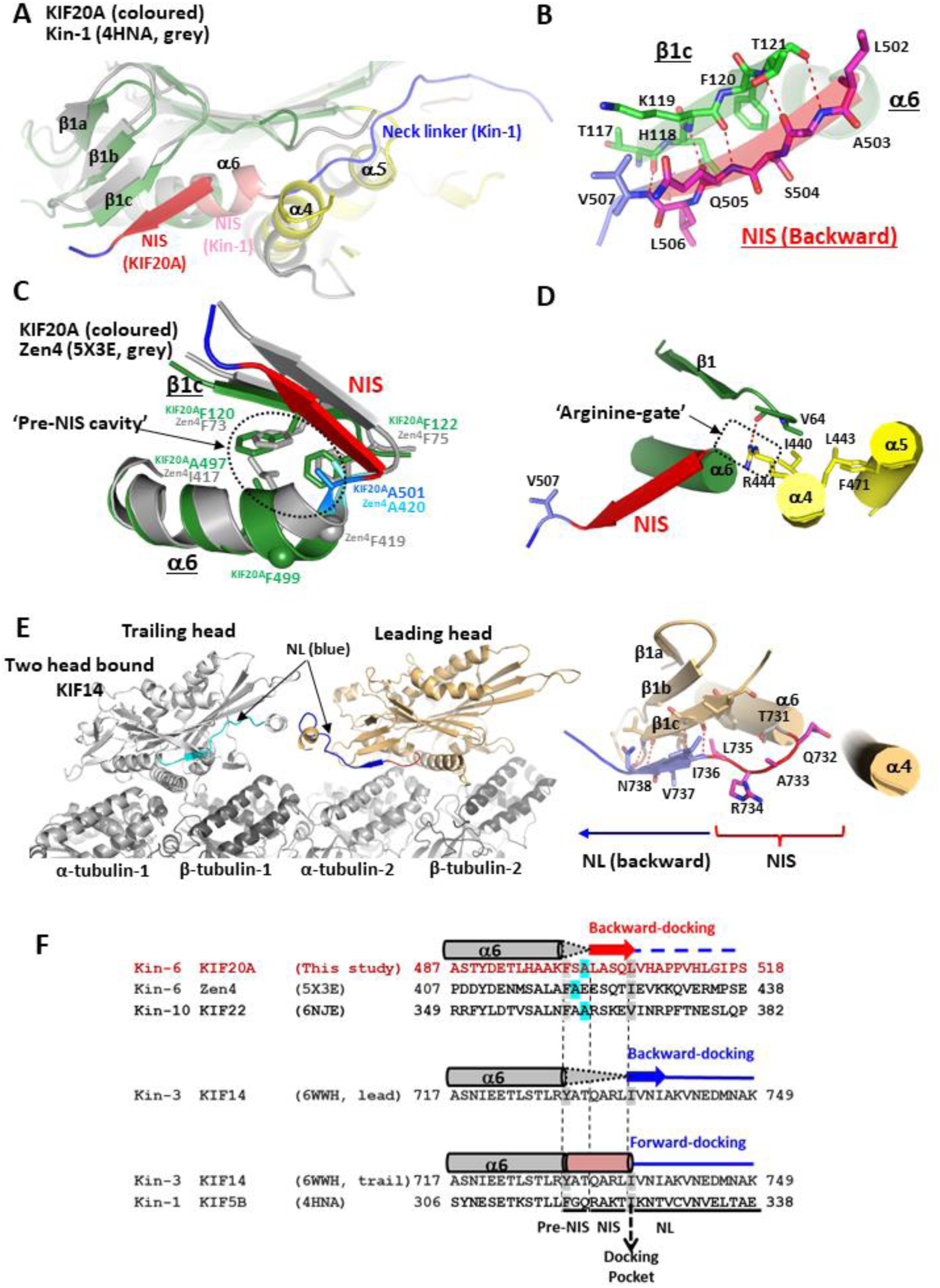
with 2 supplements. Conformation of the NL in the open state of KIF20A. **(*A*)** The minus subdomain of the KIF20A MD structure superimposed on that of the closed structure of human Kin-1 MD (PDB, 4HNA). The NIS elements (docked backward in KIF20A (red) and a part of the helix α6 in Kin-1 (salmon) are found in opposite directions. NL fragments are colored in blue. **(*B*)** β-sheet interactions between KIF20A NIS and the β1c strand, from the Minus subdomain. **(*C*)** Comparison of the α6 helix, pre-NIS and NIS found in pre-power-stroke structures. KIF20A is compared to Zen4. The KIF20A α6 helix is one residue longer than that of Zen4. In KIF20A, a serine, Ser500, is present between the last α6 residue (Phe499) and the pre-NIS residue Ala501. In Zen4, the equivalent phenylalanine is directly followed by the Pre-NIS alanine (Ala420) and its positioning in the pre-NIS cavity requires distortion of the end of the α6 helix. Following these pre-NIS residues, a similar β-sheet interaction allows the docking of the NIS element onto the β1c strand. **(*D*)** Details of the KIF20A closed docking pocket showing the side chain Arg444 (“Arg-gate”) that prevents the NIS fragment to wind and elongate the α6 helix. **(*E*)** Structure of a dimeric construct of the Kin-3 KIF14, in a two-head bound microtubule form, with the leading head exhibiting a backward orientation of the NL via a β-sheet interaction between its proximal NL residues (Ile736, Val737 and Asn738) and the β1c strand. **(*F*)** Structure-based sequence comparison of the NIS-NL fragments of KIF20A and other plus end kinesins with a similar backward docking of either their NIS or proximal NL fragment on β1c. The sequence of the equivalent fragment of the closed structure of Kin-1 is also shown, to indicate the NIS residues that extend the α6 helix (up to the “docking pocket” position) when the NL is docked forward.

In fact, recent high-resolution cryo-EM structures of the Kin-3 KIF14 (Benoit *et al*., 2021) show that when the two heads of dimeric constructs bind to microtubule, the NL of the leading head is oriented backwards through the formation of an extra strand interacting with β1c (Figure 8E and Figure 8–figure supplement 1B). However, this backward docking does not occur when only one head is bound to microtubule. Strikingly, in this case the residues involved in the additional strand are not from the NIS but from the NL (Benoit *et al*., 2021) (Figure 8F). This suggests that the tension between the two heads simultaneously attached to the microtubule can also result into a stabilized backward orientation of the NL, via a β-sheet interaction with β1c in other kinesins. This strain is not a prerequisite for the NIS docking in KIF20A, since this conformation is stable in the single head structures described here. The fact that the backward docking of the NIS is observed out of the context of a constrained microtubule-bound dimeric kinesin configuration implies that elements within the MD favour this conformation.

We find that one critical element is the pre-NIS sequence. In both KIF20A and Zen4 structures, the side chains of the NIS residues do not significantly participate in the backward-docking stabilization. In fact, in Zen4, the NIS starts one residue earlier in sequence, with the preceding α6 helix shorter by one residue, and the pre-NIS linker thus adopts a different conformation (Figure 8C). The conserved feature of these structures is the positioning of an alanine of the pre-NIS linker (^KIF20A^Ala501 and ^Zen4^Ala420) so that it occupies the same hydrophobic pocket, that we call the pre-NIS cavity, adapted for hosting only a small side chain (Ala or Gly) of the pre-NIS (Figure 8C). A similar pre-NIS alanine residue is also found in the ADP-bound Kin-10 KIF22 structure that displays a similar NIS backward-docking (Figure 8F).

By contrast, when examining the NIS sequence from other plus-end kinesins, the presence of an arginine-alanine-lysine ‘RAK’ motif (Ren *et al*., 2018) appears to favor an extended α6 helix (Figure 8–figure supplements 1A, 1C and 2). The lysine residue of this motif (^Kin-1^Lys323) lies in proximity to the ^Kin-1^Asp49 when the NIS stays helical. When conserved in plus-end kinesins, this aspartate can thus contribute to the winding and stabilization of the extended α6 helix. However, both the aspartate and the ‘RAK’ motif are missing in Kin-6s and the residues found at these positions are inadequate to interact with one another (Figure 8–figure supplement 2). The absence of this α6 helix stabilization in Kin-6s can thus favor unwinding and backward docking of the NIS. In the case of the lead head of KIF14, Thr731 cannot fit in the pre-NIS cavity and the following NIS ‘QAR’ residues found in place of the ‘RAK’ motif cannot dock on β1c (Figure 8E and F). In fact, both the pre-NIS and NIS do not wind as part of α6, but they form a disordered undocked linker. The backward docking to extend the β1a-β1c β-sheet in fact only involves NL residues via the formation of three hydrogen bonds (Figure 8E).

In the nearby α4 helix, the KIF20A Arg444 (the equivalent of the Zen4 “Arginine Gate” Arg363 (Guan *et al*., 2017) provides a large side chain that seems incompatible with the helical conformation of the last turn of the α6 helix in this open state (Figure 8D). Furthermore, Arg444 stabilizes this state by making a strong hydrogen bond with the carbonyl of Val64, which anchors the α4 helix to the MD N-terminal fragment and blocks access to the pocket that accommodates forward docking of the NL during the power stroke (Figure 8D) (Budaitis *et al*., 2019; Cao *et al*., 2017; Sindelar, 2011). Interestingly, the residue found at the ‘arginine gate’ position of the α4 helix is a conserved arginine in Kin-6s but not in other classes, in which a residue with a short aliphatic side chain (Val, Ala; Ala129 in Kin-1) or a small polar side chain (Asn312 in KIF22) is found (Figure 8–figure supplement 2). Therefore, a large (e.g. ^KIF20A^Arg444) and/or polar (e.g. ^KIF22^Asn312) residue at the “Arginine gate” position disfavors the helical conformation of the NIS, increasing its propensity to be docked backwards. Thus, the nature of the residues that can interact or clash with the pre-NIS cavity as well as those involved in stabilization of the helical form of the NIS likely contribute to tune the NIS docking/unwinding or rewinding.

In conclusion, the pre-NIS and the NIS sequences, as well as the surrounding residues involved in their stabilization differ among kinesins and likely cooperate to modulate the NIS conformation, and therefore the kinetics of transitions for the power stroke.

### The N- and C-terminal extensions contribute to KIF20A functionalities

The large extensions at both the N- and C-termini of the MD constitute two other distinct features in KIF20A (Figure 1A and Figure 1–figure supplement 1). At the C-terminus, the NL fragment (506-553), which links the MD to the dimerization domain, is ~4 times longer in KIF20A than in Kin-1 (Figure 9A). Such a long NL should preclude processive high-speed motility (Andreasson *et al*., 2015; Budaitis *et al*., 2019; Clancy *et al*., 2011; Düselder *et al*., 2012; Hackney *et al*., 2003; Ren *et al*., 2018; Shastry & Hancock, 2010; Yildiz *et al*., 2008). Likewise, the “cover-strand” (CS), i.e. the N-terminal fragment that assists the power stroke by forming the “cover-neck bundle” (Budaitis *et al*., 2019; Hwang *et al*., 2008; Khalil *et al*., 2008) corresponds to residues 55-64 in KIF20A (Figure 1–figure supplement 1), which raise questions on the role of the 1-54 fragment. We confirmed the involvement of the KIF20A N-terminal extension with a gliding motility assay, using dimeric constructs with shortened N-terminal extension 55-665 and 25-665 (Figure 9B, C and Figure 5–figure supplement 2). The gliding speed measured at 23°C produced by the 55-665 construct was lower than that produced by the 1-665 construct, implying that the N-terminal extension has a role in controlling KIF20A velocity (Figure 9B and C). The presence of the entire N-terminal extension appears necessary, since an intermediate truncation (25-665) displayed an even lower velocity (21 nm/s, Figure 9C and Figure 5–figure supplement 2A-D and Video 3). We repeated the gliding assays at 30°C to control for temperature effects and found only neglectable increases in speed (Figure 5–figure supplement 2).

**Figure 9.**
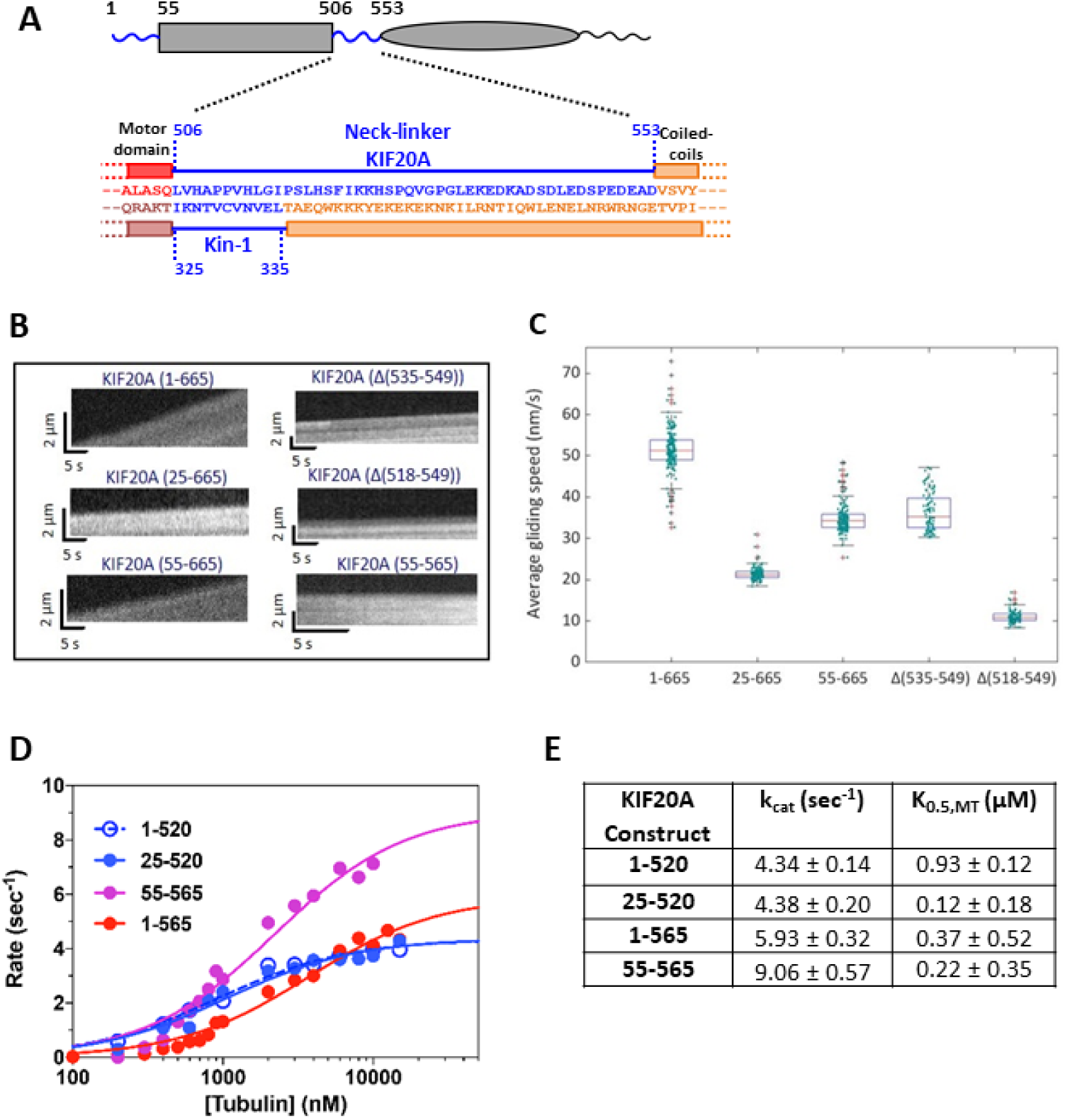
with 1 supplement. Motility and ATPase activities of different constructs of KIF20A having different truncations or deletions in the N-terminal extension or in the NL. (*A*) Schematic representation of KIF20A highlighting the long N-terminal and NL extensions (blue). The 48 residue-long sequence of KIF20A NL is compared to that of Kin-1 (11 residues). (*B*) Kymographs of TMR-microtubules propelled by the KIF20A dimeric motor constructs 1-665, 25-665, 55-665, Δ(535-549), Δ(518-549) and the monomeric 55-565 (control), respectively, at 23°C. The vertical axis of the kymographs represents microtubule displacement, the horizontal axis time. (*C*) Average gliding speeds produced by the constructs in *B*. (*D*) microtubule-stimulated ATPase for four KIF20A monomeric constructs. (*E*) Steady-state parameters extracted from *D*.

However, although we used the monomeric 1-565 construct that contains both the intact N- and C-terminal extensions for Cryo-EM (Figure 1A and Figure 2A), no density was present for the residues 1-58 and 518-565. This indicates that the NF state of KIF20A is associated with disordered N-terminal and NL extensions, even when microtubules are present. Interestingly, while no prediction can be made for the N-terminal extension from the KIF20A-microtubule complex structure, the backward directed NL (506-517) is positioned towards the microtubule minus end in such a way that it contacts residues of α-tubulin alongside the L2 loop (Figure 7). Further gliding motility assays with deletion constructs in the NL shed more lights into the functional importance of specific NL residues. The first construct lacked a stretch of 15 charged residues at the C-terminal end of the NL (Δ(535-549)), the second lacked a further 17 residues (Δ(518-549)), leaving a NL of 16 residues, slightly longer than the 10 residues NL found in transport kinesins. The construct with the shortest NL (Δ(518-549)) displayed a very low gliding velocity (10.6 ± 0.6 nm/s) (Figure 9B, 9C, Figure 5–figure supplement 2A, 2C and Video 5), while the gliding velocity produced by the Δ(535-549) deletion mutant (35.7 ± 4.4 nm/s, Figure 9B, 9C, Figure 5–figure supplement 2A1 2C and Video 4) was closer to that of the parental construct, suggesting that the charged residues at the C-terminus of the NL are not essential for the collective driving of microtubules motility. Consistently, by examining KIF20A sequences from various species, the level of conservation of the NL sequence up to residue 531 is higher than for downstream residues, indicating that within the NL, the fragment 506-531 has an evolutionary important role (Figure 9–figure supplement 1).

We also confirm the role of the N- and C-terminal extensions in modulating the kinetic properties of KIF20A with steady-state kinetics assays with different deletion constructs (Figure 9D and E). Compared to the 1-565 constructs, those with shortened NLs (1-520 and 25-520) displayed an approximately 40% reduction in k_cat_ and a 2-3 fold increase in K_0.5,MT_ (Figure 9D and E). Reducing the length of the N-terminal extension, on the other hand, increased k_cat_ by about 70% with a modest reduction in K_0.5,MT_ (Figure 9D and E). These results suggest that the N-terminal extension does not modulate the affinity of KIF20A towards microtubules. Efficient ATP hydrolysis however requires a full N-terminal extension beyond the CS as well as a much longer NL compared to other kinesins. Thus, these extensions to the MD core play essential roles in regulating enzymatic turnover in KIF20A.

### Concluding remarks

Altogether, our results highlight several structural features in KIF20A that contribute to explain why this kinesin exhibits a weak affinity for nucleotide even when microtubule is absent. This property most likely concur to the observed low velocity, when compared to effective transporters such as Kin-1 or

Kin-3. With this ability to stably reach the open NF conformation, this Kin-6 sets apart from other classes of plus-end directed kinesins and – unlike other kinesins which tightly bind ADPs in the absence of microtubule – displays a mechanism in which ADP is released before the engagement on the microtubule track (Figure 10).

**Figure 10.**
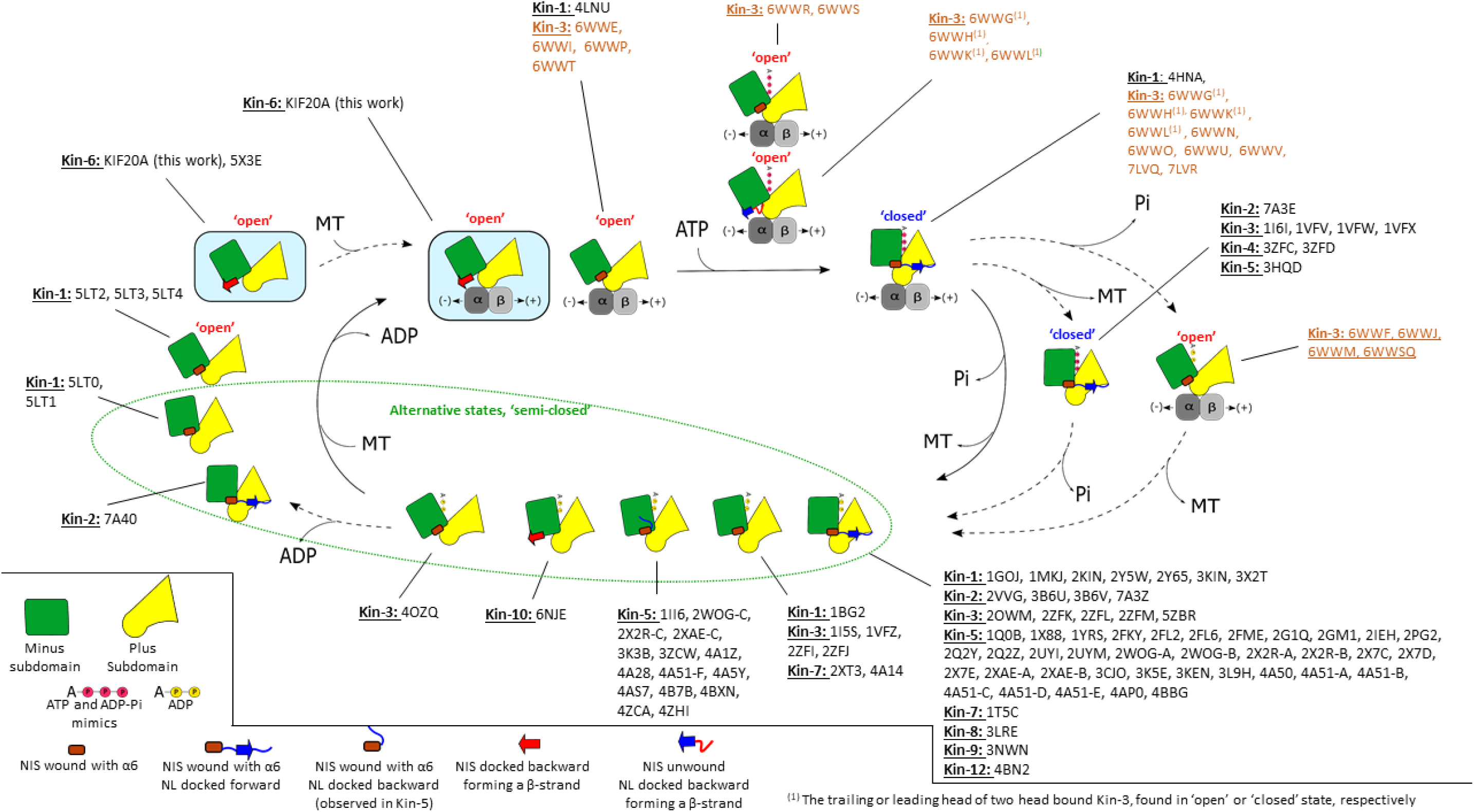
Schematic representation of the KIF20A NF conformation along a simplified generic motor cycle, showed with other states seen in different available structures of plus end kinesins. The twist of the central β-sheet is represented by different relative orientations of the minus and plus subdomains. The KIF20A structures of the current study are highlighted on a light blue background. The other crystal structures reported (PDB IDs in black) are those at higher resolution than 3.7 Å. The recent high-resolution cryo-EM structures of the microtubule-bound Kin-3 KIF14 at different nucleotide states have been included (PDB IDs in orange).

## Discussion

KIF20A is an atypical member of the kinesin superfamily with key roles during the cell cycle. The unique and long loop insertions in its MD, as well as N-terminal and NL large extensions are likely to contribute to the cellular function of this Kin-6. Our high-resolution structure of the MD – alone or bound to the microtubule – and our functional assays enlighten how these KIF20A-specific elements impact motor function. We show how a large part of the 104-residue-long KIF20A L6 loop, the L6-N fragment, strongly interacts with the plus and minus subdomains. This influences the KIF20A kinetics by affecting structural changes along the ATPase cycle and/or microtubule affinity. The L6-M moiety is disordered in our structures. This portion is likely to constitute a supplemental modulating element for this kinesin in cells by associating with cellular partners, either cargoes or regulatory proteins. Therefore, the L6 loop emerges as a major contributor to the specialization of KIF20A. Such a role of the L6 loop echoes the modulatory role for force generation in myosin splice variants that has been demonstrated in *Drosophila* muscle myosin via alternative splicing in the exon 7 (Miller *et al*., 2007). It is likely that this element alters the transitions of the motor cycle by impacting how the rearrangements can occur in the central β-sheet.

Strikingly, when compared to other plus end directed motile kinesins, KIF20A does not display a strong affinity to nucleotide. We found that this kinesin unusually stabilizes the open conformation of the nucleotide binding site in the absence of microtubule. In contrast, structural comparison indicates that the KIF20A sequence might result from necessary structural adaptation to a distinct mechanical role in which destabilization of the conformation of higher affinity for Mg-ADP compared to that observed for other processive kinesins such as Kin-1 would be an advantage. Analysis of the structure and the specific sequence of this kinesin indicates that the unusually large L6 can contribute to this feature via a direct influence of the L6-N region on the twist of the central β-sheet. In addition, the C-terminal NL and the preceding NIS region further stabilize the open state by their interaction with the MD in the backwards position. As a result, KIF20A exhibits an unusual coupling between microtubules and nucleotide binding events although microtubules do activate its ATPase activity. These structural characteristics are likely to render this kinesin particularly slow as revealed by our gliding assays.

In this open state, the NIS portion of KIF20A is unwound from the α6 helix and docks on the β-strand β1c to orient the NL towards the minus-end of microtubule, like in Zen4 (Guan *et al*., 2017) or in the Kin-10 KIF22 (6NJE, Walker *et al*., unpublished). In fact, several pieces of earlier evidence support a backward orientation of the NL in some kinesins in the NF state (Goulet *et al*., 2014; Guan *et al*., 2017; Mickolajczyk *et al*., 2019; Rice *et al*., 1999; Sablin & Fletterick, 2004). As demonstrated by the recent high resolution cryo-EM study of the Kin-3 KIF14, such a backward docking is indeed found for the lead head in the two-head-bound state, albeit not in the NF state. However, we find here that the NIS (the preceding sequence) can play a critical role to favour this backward direction of the NL on a single head and without binding to microtubule. Note that the backward docking proposed from low resolution cryo-EM maps for Kin-5 disposes the NL on top of the β1a-β1c β-sheet, guided by a class specific sequence conservation (Goulet *et al*., 2014; Turner *et al*., 2001), (Figure 8–figure supplements 1 and 2C). The docking of the NIS fragment, rather than the NL *per se* (as seen in KIF14) is clearly unaccustomed among kinesins and is a characteristic of only a few kinesins that possess a residue of small side chain in the NIS. Together with the elements that control the NBS opening, the sequence elements that guide the (un)winding of the NIS and its backward docking constitute another leverage to regulate the motor mechanism and can contribute to the specific kinetics and motility properties of KIF20A.

We finally found that albeit largely disordered in our structures of the NF state, extensions at both N- and C-terminus of the MD are involved in KIF20A functionalities. Our kinetic studies indicate that in KIF20A, these elements must be much longer than in other kinesins to retain normal enzymatic properties. The mechanism beyond this property remains to be investigated in detail. It is possible that the N-terminal extension expands the cover strand to assist the NL movement during the power stroke. Future structural studies of other nucleotide states will be of high interest to highlight the role of these extensions. The current model for kinesin stepping attribute an important role of the length of the NL for processivity (Andreasson *et al*., 2015; Yildiz *et al*., 2008) and imply that the long KIF20A NL impedes processive motility. Indeed, tail-less versions of KIF20A confirm that the dimerization domain alone is not sufficient for this motor to be robustly processive (Figure 5E, 5F, 5H), which is consequently unable to transport cargoes in cell (Poulos *et al*., 2022). Interestingly, however, another study found full-length KIF20A to be processive and characterized the influence of cargoes on the run-length (Adriaans *et al*., 2020). How the tail fragment and physiological KIF20A partners influence KIF20A motility constitutes another fascinating area for future research on this kinesin. These include the proteins that are involved in KIF20A regulation and modulate its activities along the cell cycle.

For example, during the metaphase/anaphase transition in mitosis, KIF20A, bound to the CPC, acts as a transporter (Adriaans *et al*., 2020). Plk1 phosphorylates KIF20A in its NL (Ser527), which is required for Plk1 to bind to KIF20A and reach its proper localization (Neef *et al*., 2003). In interphase, KIF20A promotes membrane scission to form RAB6 vesicles at Golgi hotspots in association with myosin-2 and RAB6. At these sites, KIF20A might favor the formation of a microtubule network linked to contractile actomyosin rather than acting as a transporter since its depletion results in the appearance of elongated tubules growing from RAB6 hot spots (Miserey-Lenkei *et al*., 2017). As is the case with many other macromolecular machines, important puzzles about the complex functions of the KIF20A kinesin in the full cellular context thus await further investigation, after basic structural and functional features have been discovered.

## Materials and Methods

### Protein expression and purification

The production procedure for KIF20A constructs containing the MD was derived from the preparation method for the 25-520 KIF20A construct previously studied by Cryo-EM (Atherton *et al*., 2017). Briefly, all these constructs were cloned into pET28b with a C-terminal 6His tag and were expressed in NiCO21 (DE3) *E. coli* cells (New England BioLabs, Evry, France) by induction with 0.2 mM IPTG overnight at 20°C. Cells were harvested by centrifugation at 4500×g for 20 min and lysed by sonication in lysis buffer (50 mM HEPES, pH 7.5, 500 mM NaCl, 40 mM Imidazole, 0.5 mM TCEP, 0.1 mM ATP, 5 mM MgCl2, 5% Glycerol and Complete EDTA-free antiprotease cocktail (Roche, Boulogne-Billancourt, France)). The lysate was clarified at 20,000×g for 45 min at 4°C, and the impurities that were tagged by Chitin Binding Domain were extracted on a gravity-flow column using 5 mL Chitin resin (New England BioLabs, Evry, France). The target proteins were purified with HisTrap HP columns (GE Life Sciences, Velizy-Villacoublay, France). Further purification was achieved by size-exclusion chromatography in buffer containing 20 mM HEPES, pH 7.5, 150 mM NaCl, 1 mM TCEP, 5 mM MgCl_2_, 0.1 mM ATP. Fractions containing purified KIF20A protein were pooled and concentrated to the desired concentrations, flash-frozen in liquid nitrogen and stored in −80°C until use. All the monomeric constructs (1-520, 25-520, 55-510, 55-565, 1-565) were concentrated up to ~20 mg/mL and the dimeric constructs (1-665, 25-665 and 55-665), to ~5 mg/mL. To analyze the role of the neck linker for motor properties *in vitro*, two variants of the 55-665 dimeric motor were designed, lacking residues 518-549 or 535-549, respectively. For the 1-565 construct used for Cryo-EM, the final size-exclusion chromatography was performed with a buffer composed of 80 mM PIPES pH 6.8, 75 mM KCl, 2 mM de MgCl2, TCEP 1mM, 1mM EGTA, 0.1 mM ATP.

Verification of the monomeric or dimeric state of those constructs was obtained by SEC-MALS experiments with the injection of 5 mg/mL of each protein into a Superdex 200 10/30 Increase column (GE Life Sciences, Velizy-Villacoublay, France) mounted on a HPLC Ultimate 3000 (Thermo Scientific, Waltham, MA USA) coupled to a miniDAWN Treos and an Optilab T-rEX (Wyatt Technology, Santa Barbara, CA, USA) for MALS and refractive index measurements. Data were analyzed using ASTRA 6 software (Wyatt Technology, Santa Barbara, CA, USA).

To ensure the NF state of the 55-510 construct used for crystallization, a batch was produced with the omission of ADP in all buffers. For crystallographic structure solution, a selenomethionine-derivative of the 55-510 construct was produced using the SeMet kit from Molecular Dimensions (Sheffield, UK) using the instructions from the provider.

For control experiments, including gliding assays and single-molecule motility assays we used a truncated dimeric Kin-1 construct from *Neurospora crassa* (NK433) (Lakämper *et al*., 2010), a Kin-3 construct from *Caenorhabditis elegans* (UNC-104 U653) (Klopfenstein *et al*., 2002; Tomishige *et al*., 2002) and a chimeric Kin-5/Kin-1 construct (chimeric Eg5Kin-NL-18) (Düselder *et al*., 2012). The motors were expressed using the E. coli BL21 (DE3) strain. NK433 and Eg5Kin-NL-18 were expressed and purified essentially as described previously (Düselder *et al*., 2012). Briefly, when bacterial growth reached exponential phase, protein expression was induced with 0.1 mM IPTG overnight at 20°C. Cells were harvested and sonicated as described above, in 20 mM NaPO_4_, pH 7.4, 1 mM MgCl_2_, 10 mM β-mercaptoethanol, 0.1 mM ATP. The target proteins were first purified by Ni-NTA affinity chromatography, then buffer-exchanged by overnight dialysis in BRB80 buffer (80 mM PIPES, pH 6.8, 1 mM MgCl_2_, 1 mM EGTA) supplemented with 0.1 mM ATP and 1 mM DTT. UNC-104 U653 was further purified by ion-exchange chromatography and was eluted in a ratio of 70 % buffer A (25 mM PIPES, pH 6.8, 2 mM MgCl_2_, 0.1 mM EGTA, 0.1 mM ATP, 1 mM DTT) and 30 % buffer B (25 mM PIPES, pH 6.8, 2 mM MgCl_2_, 0.1 mM EGTA, 250 mM NaCl, 0.1 mM ATP, 1 mM DTT). Aliquots were then snap frozen in liquid nitrogen and stored at −80°C.

### Microtubule preparation

For motility assays, all reagents were obtained from Sigma-Aldrich (Hamburg, Germany) unless otherwise noted. Tubulin was purified from pig brain and labeled with Tetramethyl-rhodamine (TMR) as described (Hyman *et al*., 1991; Williams & Lee, 1982) and stored at −80°C after flash freezing in liquid N_2_. Partially TMR-labeled microtubules were polymerized from 1.7 μM 1.6:1 Tetramethyl-rhodamine labeled tubulin and 33 μM unlabeled tubulin in the presence of 1 mM GTP, 4 mM MgCl_2_ in BRB80 buffer (80 mM Pipes/KOH, pH 6.8, 1 mM MgCl_2_, and 1 mM EGTA), incubated for 20 min at 37°C. The microtubules were stabilized in BRB80 containing 10 μM Paclitaxel (Taxol®).

For cryoEM experiments, microtubules were prepared from porcine brain tubulin (Cytoskeleton, Inc. CO). Tubulin lyophilized pellets were resuspended in BRB80 (80 mM K-PIPES, 1 mM MgCl_2_, 1 mM EGTA, pH 6.8) to 5 mg/ml and spun at 313,000 × g before polymerization to eliminate aggregates. microtubule polymerization was done in conditions to enrich the number of microtubules with 15 protofilaments (Wilson-Kubalek *et al*., 2016) as follows. The clarified resuspended tubulin solution was supplemented with 2 mM GTP, 4mM MgCl_2_, 12%(v/v) DMSO and incubated 40 min at 37 °C. An aliquot of stock Paclitaxel (Taxol®) solution (2 mM in DMSO) was added for a final Paclitaxel concentration of 250 µM and incubated for another 40 min at 37 °C. The microtubules were then spun at 15,500 × g, 25 °C and the pellet resuspended in BRB80 with 20 μM paclitaxel.

### Determination of the KIF20A 55-510 NF structure by X-ray crystallography

#### Crystallization

The 55-510 construct (20 mg/mL) led to agglomerates of crystals in vapor diffusion setups at 17°C, in 2 M NaCl, 0.5 M (NH_4_)_2_SO_4_, 0.1 M Bis-Tris pH 5.5, by mixing 1 µL of protein sample with an equal volume of precipitant. The initial crystals were optimized by seeding in 1.8 M NaCl, 0.5 M (NH_4_)_2_SO_4_, 0.1 M Bis-Tris, pH 5.5. Crystals were then harvested, cryo-protected with the above buffer supplemented with 20 % glycerol and flash-frozen in liquid nitrogen for diffraction data collection.

#### Diffraction data collection and processing, structure determination and refinement

Diffraction data were collected at the Proxima-1 beamline of the synchrotron SOLEIL (Saint-Aubin, France). Complete datasets were collected from individual crystals under a cryogenic stream at 100 K. All datasets were collected with a wavelength of 0.98 Å and processed using the automated pipeline AutoProc (Vonrhein *et al*., 2011) that executes XDS (Kabsch, 2010), Pointless (P. Evans, 2006) and Aimless (P. R. Evans & Murshudov, 2013) of the CCP4 suite (Winn *et al*., 2011). Se-SAD phasing was first performed using AutoSharp (Vonrhein *et al*., 2011), with a crystal grown in presence of 2 mM Mg^2+^.ADP. No ADP was visible in the nucleotide binding site of the resulting model, which was thus refined against another native crystal dataset obtained in nucleotide-free conditions. This model was extended by iteratively using Buster (Bricogne *et al*., 2020) and Coot (Emsley *et al*., 2010). The refinement statistics of the final model (2.7 Å resolution) are shown in Supplementary file 1.

### Determination of the KIF20A 1-565 NF structure in complex with microtubule by cryo-EM

#### Sample preparation

Four µL of 6 μM microtubules BRB80 plus 20 μM paclitaxel were layered onto a (UltrAuFoil R1.2/1.3 300 mesh) plasma cleaned gold grid just before use (Gatan Solarus plasma cleaner, at 15 W for 6 s in a 75% argon/25% oxygen atmosphere), the microtubules were incubated 1 min at room temperature and then the excess liquid removed from the grid using a Whatman #1 paper. Four µl of the 40 μM Kif20A in BRB40 supplemented with 20 μM paclitaxel and 20 × 10^−3^ unit of potato apyrase (Sigma-Aldrich, MO), were then applied onto the EM grid and incubated for 1 min at room temperature. The grid was mounted into a Vitrobot apparatus (FEI-ThermoFisher MA), incubated 1 min at room temperature and plunged frozen into liquid ethane. Vitrobot settings: 100% humidity, 3 s blotting with Whatman #1 paper and −2 mm offset. Grids were stored in liquid nitrogen until imaging in a cryo-electron microscope.

#### Cryo-EM data collection

Data were collected at 300 kV on a Titan Krios microscope equipped with a K3 summit detector. Acquisition was controlled using Leginon (Suloway *et al*., 2005) with the image-shift protocol (to target ~30 holes per stage shift) and partial correction for coma induced by beam tilt (Glaeser *et al*., 2011). The pixel size was 0.844 Å/pixel and the cumulated dose of 56 e^−^/Å^2^ (Supplementary file 2). The exposures were fractionated on 40 movie frames.

#### Processing of the cryo-EM datasets of microtubule-kinesin complexes

The processing was done as previously described (Benoit *et al*., 2021). Movie frames were aligned with Relion generating both dose-weighted and non-dose-weighted sums. Contrast transfer function (CTF) parameters per micrographs were estimated with Gctf (K. Zhang, 2016) on aligned and non-dose-weighted movie averages.

Helical reconstruction on 15R microtubule was performed using a helical-single-particle 3D analysis workflow in Frealign (Grigorieff, 2016), as described previously (Benoit *et al*., 2018, 2021) with 664 pixels box size, with each filament contributing only to one of the two half dataset. Per-particle CTF refinement was performed with FrealignX (Grant *et al*., 2018).

To select for tubulins bound to kinesin motors and to improve the resolution of the kinesin-tubulin complexes, the procedure HSARC (Benoit *et al*., 2021) was used for these one-head-bound states. The procedure follows these steps:

1. Relion helical refinement. The two independent Frealign helical refined half datasets were subjected to a single helical refinement in Relion 3.1 (Zivanov *et al*., 2018) where each dataset was assigned to a distinct half-set and using as priors the Euler angle values determined in the helical-single-particle 3D reconstruction (initial resolution: 8 Å, sigma on Euler angles sigma_ang: 1.5, no helical parameter search).
2. Asymmetric refinement with partial signal subtraction. An atomic model of a kinesin-tubulin complex was used to generate two soft masks using EMAN pdb2mrc and relion_mask_create (low-pass filtration: 30 Å, initial threshold: 0.05, extension: 14 pixels, soft edge: 8 pixels). One mask (mask_full_) was generated from a kinesin model bound to one tubulin dimer and two longitudinally flanking tubulin subunits while the other mask (mask_kinesin_) was generated with only the kinesin coordinates. The helical dataset alignment file was symmetry expanded using the 15R microtubule symmetry of the dataset. Partial signal subtraction was then performed using mask_full_ to retain the signal within that mask. During this procedure, images were re-centered on the projections of 3D coordinates of the center of mass of mask_full_ (C_M_) using a 416 pixels box size. The partially signal subtracted dataset was then used in a Relion 3D refinement procedure using as priors the Euler angle values determined from the Relion helical refinement and the symmetry expansion procedure (initial resolution: 8 Å, sigma_ang: 5, offset range corresponding to 5 Å, healpix_order and auto_local_healpix_order set to 5). The CTF of each particle was corrected to account for their different position along the optical axis.
3. 3D classification of the kinesin signal. A second partial signal subtraction procedure identical to the first one but using mask_kinesin_ and with particles re-centered on the projections of C_M_ was performed to subtract all but the kinesin signal. The images obtained were resampled to 3.5 Å/pixel in 100 pixel boxes and the 3D refinement from step 2 was used to update the Euler angles and shifts of all particles. A 3D focused classification without images alignment and using a mask for the kinesin generated like mask_kinesin_ was then performed on the resampled dataset (8 classes, tau2_fudge: 4, padding: 2, iterations: 175). That classification lead to one class with a kinesin visible (12.8% of the dataset). As the decoration was low and the kinesin density heterogeneous, another round of 3D classification in 8 classes with the same parameters was performed on the subset of data corresponding to that decorated class. This generated 2 classes with a kinesin having some secondary structure visible, the rest being at too low resolution on the kinesin part for proper assessment. So, the decorated subset of the data was used as input in a 3D classification with only 2 classes. The final 2 classes, class-1 and class-2 comprised 66% and 33% of the input data respectively (8.6% and 4.2% of the full dataset).
4. 3D reconstructions with original images (not signal subtracted). To avoid potential artifacts introduced by the signal subtraction procedure, final 3D reconstructions were obtained using relion_reconstruct on the original image-particles without signal subtraction.

To obtain a final locally filtered and locally sharpened map for the higher resolution class-1, post-processing of the pair of unfiltered and unsharpened half maps was performed as follows. One of the two unfiltered half-map was low-pass-filtered to 15 Å and the minimal threshold value that does not show noise around the microtubule fragment was used to generate a mask with relion_mask_create (low-pass filtration: 15 Å, extension: 10 pixels, soft edge: 10 pixels). This soft mask was used in blocres (Heymann & Belnap, 2007) on 12-pixel size boxes to obtain initial local resolution estimates. The merged map was locally filtered by blocfilt (Heymann & Belnap, 2007) using blocres local resolution estimates and then used for local low-pass filtration and local sharpening in localdeblur (Ramírez-Aportela *et al*., 2020) with resolution search up to 25 Å. The localdeblur program converged to a filtration appropriate for the tubulin part of the map but over-sharpened for the kinesin part. The maps at every localdeblur cycle were saved and the map with better filtration for the kinesin part area was selected with the aim of better resolving the kinesin loops. For the lower resolution class-2, the map was postprocessed in Relion, applying a bfactor of −80 Å^2^ and low pass filtered at 4 Å.

#### Cryo-EM resolution estimation

The final resolutions for each cryo-EM class average reconstruction were estimated from FSC curves generated with Relion 3.1 postprocess (FSC_0.143_ criteria, Figure 2–figure supplement 2). To estimate the overall resolution, these curves were computed from the two independently refined half maps (gold standard) using soft masks that isolate a single asymmetric unit containing a kinesin and a tubulin dimer. The soft masks were created with Relion 3.1 relion_mask_create (for microtubule datasets: low pass filtration: 15 Å, threshold: 0.1, extension: 2 pixels, soft edge: 5 pixels) applied on the correctly positioned EMAN pdb2mrc density map generated with the coordinates of the respective refined atomic models. FSC curves for the tubulin or kinesin parts of the maps were generated similarly using the corresponding subset of the PDB model to mask only a kinesin or a tubulin dimer (Supplementary file 2).

### Steady state ATPase assays and pre-steady state kinetics

ATPase assays were performed using the EnzChek phosphate assay kit (Invitrogen) with motor concentrations in the 5 to 50 nM range and 25nM to 2 µM polymerized tubulin. ATPase assays were performed in 25 mM HEPES, 50 mM potassium acetate, 5 mM magnesium acetate, 1 mM EGTA, and 1 mM DTT, pH 7.5. k_cat_ and K_0.5,MT_ were calculated from fitting the Michaelis-Menten equation to ATPase data taken at a >10-fold excess of microtubules.

Pre-steady-state kinetics of 2’ deoxy 3’ MANT-ATP (2’dmT) and 2’ deoxy 3’ MANT-ADP (2’dmD) binding, nucleotide-dependent microtubule release and Pi release were performed on a KinTex SF-2004 stopped-flow apparatus as previously described (Muretta *et al*., 2013).

Binding of the fluorescent nucleotide analogues 2’ deoxy 3’ MANT-ADP (2’dmD) and 2’ deoxy 3’ MANT-ATP (2’dmT) to 4:1 microtubules:motor complex were measured by mixing with an excess of fluorescent nucleotide in a KinTek F-300X stopped flow at 20°C, with instrument dead time of 1.2 milliseconds. Samples were rendered nucleotide free prior to mixing by incubating for 20 min with 0.2 U/ml apyrase (Type VII, Sigma Aldrich, St Louis, MO). Fluorescence enhancement of the MANT-fluorophore was monitored by energy transfer from vicinal tryptophans by exciting at 295 nm and monitoring 90° from the incident beam through a 450 nm broad bandpass filter (Omega Optical, Brattleboro, VT). Data were subjected to linear least squares fitting.

### Gliding motility assays

Gliding motility assays were performed as described previously (Lakämper *et al*., 2010) at room temperature (23°C) or at 30°C. 0.5 – 1.0 μM motor solutions were filled into assay chambers (KIF20A constructs, Kin-1 and Kin-3) and were allowed to specifically bind to the coverslips of the assay chambers pre-treated with protein G and an Anti-His antibody. Chambers were made from 22×24 mm cleaned coverslips spaced from a 75×25 mm cleaned microscope slides (Figure 5–figure supplement 1) using double-sided tape of ~100 μm thickness (Tesa SE, Hamburg, Germany). After motor attachment, chambers were filled with 10 μL tetramethyl-rhodamine (TMR)-labeled microtubules (~0.4 μM tubulin dimers in anti-bleach buffer containing 2 mM ATP). Anti-bleach buffer contained 80 μg/mL catalase, 100 μg/mL glucose oxidase, 10 mM D-glucose and 10 μM Paclitaxel, in BRB80 buffer.

Motility was observed in an inverted fluorescence microscope (Axiovert 200, Carl Zeiss, Jena, Germany) using a Zeiss EC Plan-Neofluar 100x 1.3 NA oil immersion objective. Images were recorded with a CCD camera (Photometrics CoolSnap EZ; Roper Scientific GmbH, Ottobrunn, Germany). Data were acquired with WinSpec software (Princeton Instruments, Princeton, NJ, USA). Further analysis and statistical calculations were performed with MATLAB (The MathWorks, Natick, MA, USA) and Fiji/ImageJ (NIH).

### Single-molecule motility assays

Coverslips were cleaned using 0.5 mM KOH prior to silanization with a positively charged silane, 3-[2-(2-aminoethylamino)-ethylamino] propyl-trimethoxysilane (DETA) (Sigma-Aldrich, Hamburg, Germany) for microtubule immobilization. Microtubules were incubated on the treated coverslips for 10 min followed by 10 min incubation with 0.1 mg/ml Casein (Sigma-Aldrich) in BRB80 buffer. The chamber was then washed with 10 μL of assay buffer (BRB80 buffer + anti-bleaching agents) and then filled with 10 µL of 20-50 nM of GFP-tagged motor proteins (KIF20A 1-665 and Kin-5) in assay buffer containing 2 mM ATP. Fluorescence was observed in a custom-built total internal reflection fluorescence (TIRF) microscope described previously (Fish, 2009) using an oil-immersion objective (SFluor 100x, NA 1.49, Nikon Instruments Europe BV, Amsterdam, Netherlands). Images were recorded with a CCD camera (Ixon Ultra, Andor Technology - OXFORD Instruments, Oxford, UK). Videos were recorded at 10 frames per second and were analyzed for motor speeds using kymographs generated with Solis software from Andor. Statistical analysis of the data was performed with MATLAB (The MathWorks) and Fiji/ImageJ (NIH). All experiments were performed at room temperature (23°C).

The dwell times of GFP-tagged motor proteins in the regions selected for kymographs were measured from the kymographs that were generated from unprocessed image sequences using ImageJ. Start and end points of individual dwell events were determined from kymograph intensity profiles (Figure 5 B and C) using a threshold of 10% of the maximal intensity. Note that brief appearances of fluorescent proteins can also occur due to diffusion in and out of the TIRF detection zone of about 200 nm depth near the substrate surface.

### Small angle X-Ray scattering

SAXS data were collected on the SWING beamline (synchrotron SOLEIL, France). Purified Kif20A 1-565 was incubated with either 5 mM AMPPNP, ADP or in NF conditions for 15min on ice, and then all samples were centrifuged at 20,000 × g for 10 min at 4 °C prior to the analysis. 45 μl of the protein at 4.43 mg ml^−1^ (70.4 µM) was injected in SEC column S200i 5/150 (GE Life Sciences, Velizy-Villacoublay, France) with a buffer containing 20mM Hepes pH 7,5, 1mM DTT, 150mM NaCl, 5mM MgCl2 and supplemented with 5mM of nucleotide when appropriate. All datasets were processed using the Foxtrot software (version 3.5.9) (Xenocs Soleil Synchrotron, Sassenage, France) (Tencé-Girault *et al*., 2019), then Rg value calculations were performed using the Primus software (Konarev *et al*., 2003).

## Data availability

The atomic models are available in the PDB, www.pdb.org, under accession numbers PDB 8BJS for the KIF20A-MD-NF X-ray structure (Supplementary file 1), and 8F1A and 8F18 for the cryo-EM structure (class-1 and class-2, respectively). The final cryo-EM maps together with the corresponding half maps, masks used for resolution estimation, masks used in the partial signal subtraction for the microtubule datasets, and the FSC curves are deposited in the Electron Microscopy Data Bank, under accession numbers EMD-28789 and EMD-28787 (Supplementary file 2).

## Acknowledgments

We thank I-Mei Yu for preliminary experiments performed for this project. We are grateful to the beamline scientists of PX1 (SOLEIL synchrotron) for excellent support and we thank Margaret A. Titus and members of the Structural Motility group together with Bruno Goud, Phong Thanh Tran and Stéphanie Miserey-Lenkei for critical reading of the manuscript. We thank Biokinesis, Inc. for financial support. This work was further supported by NIH NS073610-01, GM113164 (H.S.); GM130556, U54CA210190 to SSR, the European Research Council under the European Union’s Seventh Framework Programme (FP7/2007–2013)/ERC grant agreement no. 340528 to CFS; ANR/DFG project: ANR-15-CE13-0017-01 and DFG-AZ:SCHM 799/6-1 to AH and CFS. Cryo-EM data collection was performed at the Simons Electron Microscopy Center and National Resource for Automated Molecular Microscopy located at the New York Structural Biology Center, supported by grants from the Simons Foundation (SF349247), NYSTAR, and the NIH National Institute of General Medical Sciences (GM103310) with additional support from Agouron Institute (F00316) and NIH (OD019994). AH was supported by an IRP grant from CNRS, la Ligue contre le Cancer Comité de Paris RS20/75-D, ARC 2016-1-PLBIO-10-ICR-1, FRM DCM20181039553 and INCA 2016-1-PLBIO-10-ICR-1. The AH team is part of the Labex Cell(n)Scale (ANR-11-LABX-0038), which is part of the IDEX PSL (ANR-10-IDEX-0001-02).

## Supplementary Information

**Figure 1–figure supplement 1.**
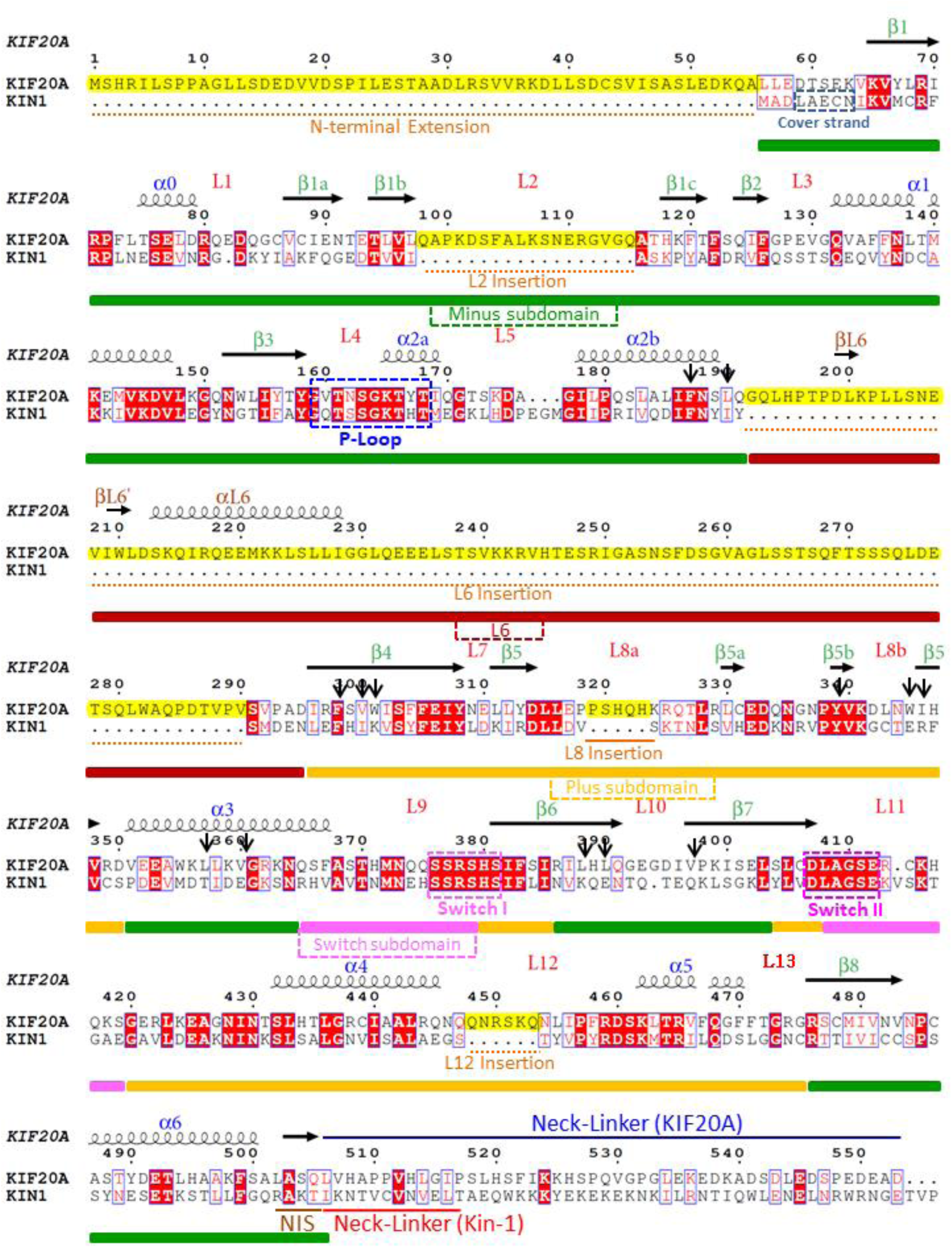
Annotated sequence alignment of the motor domains of the mouse KIF20A, and the human kinesin-1 (KIF5B) as a reference to delineate loop insertions. The plus, minus and switch subdomains (Benoit *et al*., 2021) are depicted under alignment. The MD loop insertions are highlighted in yellow, as well as the KIF20A-specific sequence at its N-terminus (N-terminal extension). Other key elements of the motor are indicated; these are the P-loop, Switch I and Switch II motives, as well as the NIS elements discussed in this study. At the N-terminus, the cover strand that forms the cover-neck bundle in the Kin-1 post-power-stroke state is also indicated (Budaitis *et al*., 2019; Hwang *et al*., 2008). Hydrophobic residues of the MD core, involved in the stabilization of the L6 insert, are indicated with an arrow. Examination of 100 non redundant kinesin sequences from various species (from yeast to human) show that 70 % of these residues are hydrophilic for kinesins that do not have a large L6 insert. The KIF20A secondary structure annotations follow an established nomenclature (Kull *et al*., 1996). This alignment was generated using ESPript 3.0 (Robert & Gouet, 2014). The KIF20A and Kin-1 (KIF5B) sequences correspond to the UniProtKB accession numbers P97329 and P33176, respectively.

**Figure 1–figure supplement 2.**
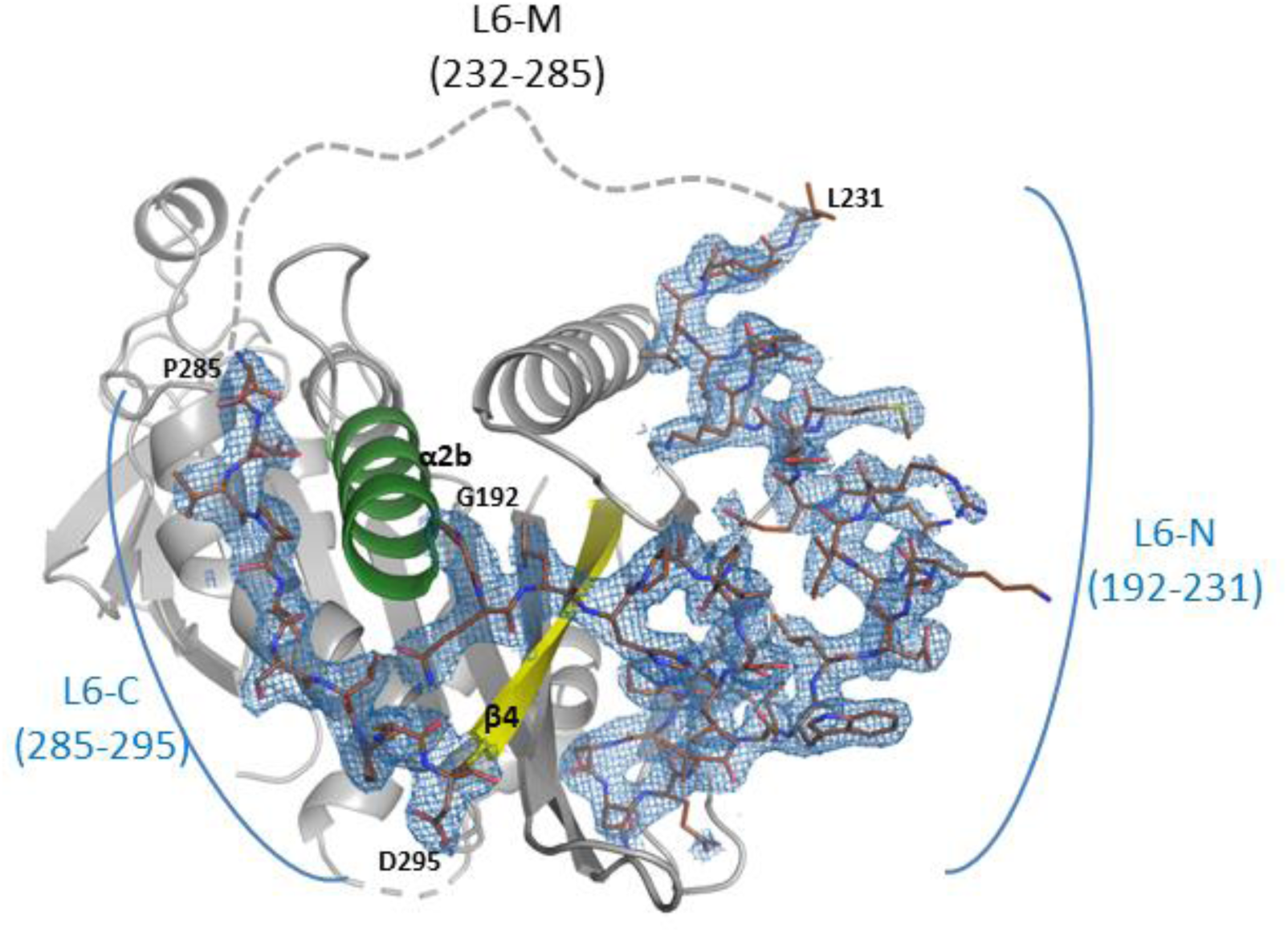
Density maps of regions of KIF20A, at 2.7 Å resolution (Crystal structure). Electron density (ED) maps at 2.7 Å resolution of areas in the vicinity of the L6 fragment 192-295 of the crystal structure (contour levels: 2mFo-DFc 1.0 rmsd).

**Figure 2–figure supplement 1.**
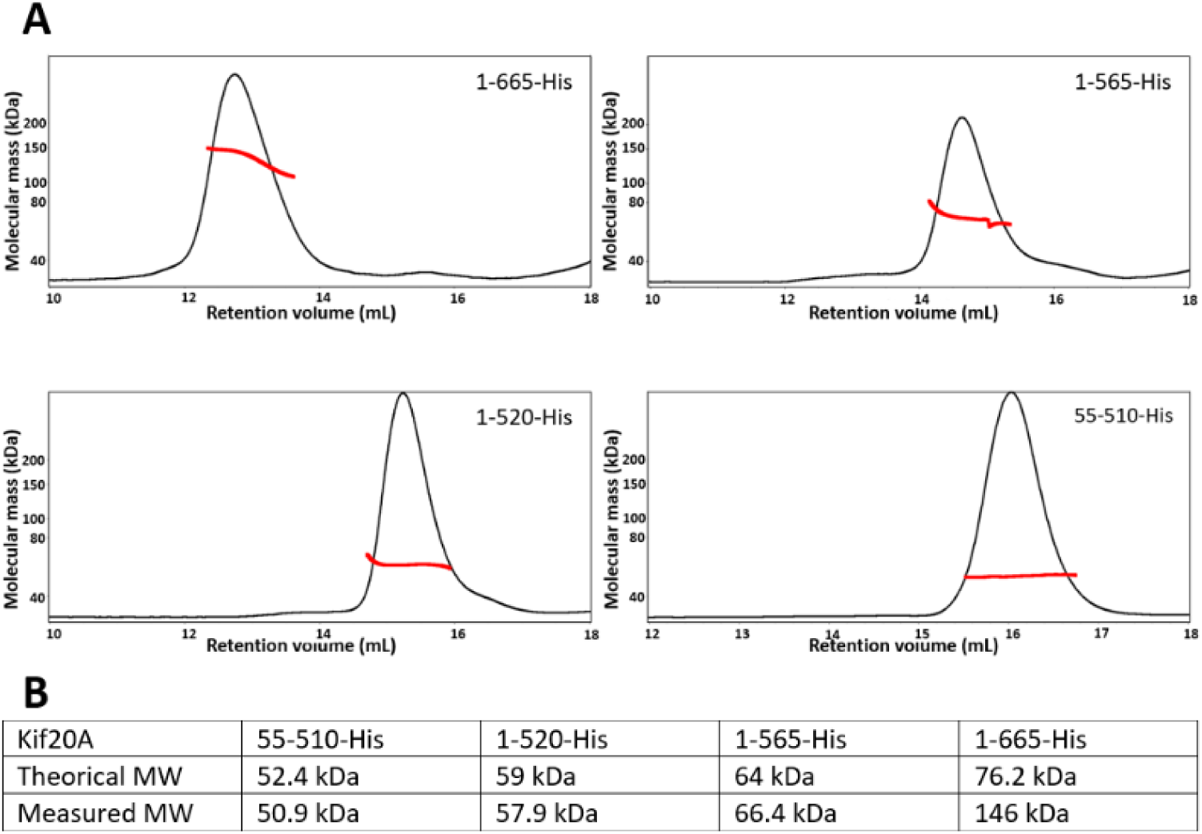
MALS experiments of the constructs used in this study. **(*A*)** Mass measured for each construct according to its retention volume. **(*B*)** Comparison of the theoretical and measured masses for each construct reflecting the oligomeric state in solution.

**Figure 2–figure supplement 2.**
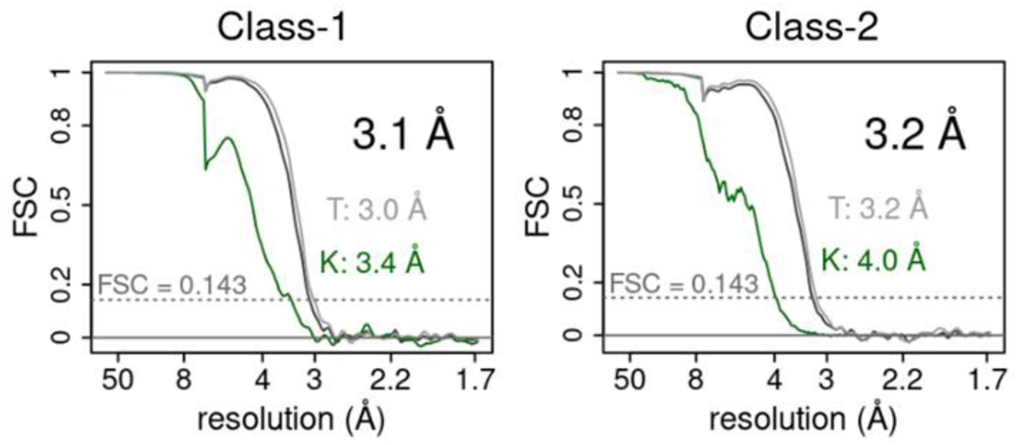
Resolution estimation of the cryo-EM maps. FSCs curves for the 2 classes solved by cryo-EM are plotted. Overall FSC in black, tubulin part FSC in grey and kinesin part FSC in green. Resolution values (FSC at 0.143) for the overall, tubulin (T) and kinesin (K) parts are indicated. Half maps and masks used to generate the FSC curves are deposited in the EMDB (accession numbers in Supplementary file 2).

**Figure 3–figure supplement 1.**
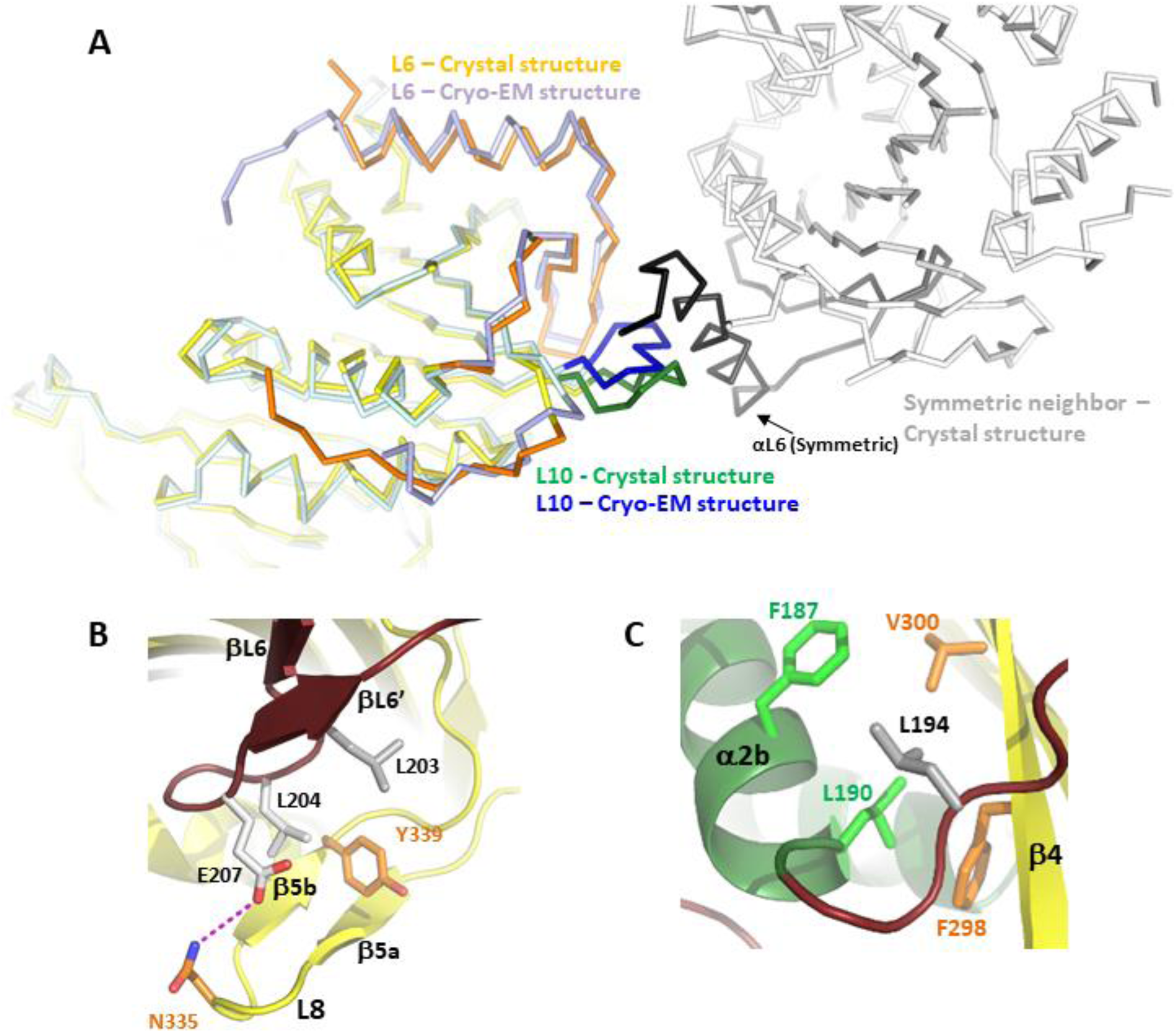
The KIF20A L6 loop: structure and interactions with the core MD. **(*A*)** Overlay between the X-ray and cryo-EM structures of KIF20A MD showing a crystal packing contact between a neighboring αL6 fragment and the L10 loop, which affects the conformation of the loop L6. As a result, the L6-N fragment is slightly displaced as compared to the cryo-EM structure (average displacement of 1.39 Å for L6-N Cα atoms, with a maximum of 2.4 Å for Lys214). **(*B*)** Contacts between the βL6-βL6’ and β5a-β5b hairpins of the L8 microtubule-binding loop illustrate how L6 could influence the interactions with the microtubule. Indeed, these interactions provide constraints to this microtubule-binding loop, not present in other kinesins, and therefore are likely to indirectly contribute to microtubule-binding mode or microtubule-binding affinity. **(*C*)** Influence of the KIF20A L6 insert on the interface between the Minus (green) and Plus (yellow) subdomains, with Leu194 inserted in a hydrophobic pocket at the subdomain interface (Phe298 and Val300 in the Plus subdomain and Phe187 and Leu190 in the Minus subdomain).

**Figure 4–figure supplement 1.**
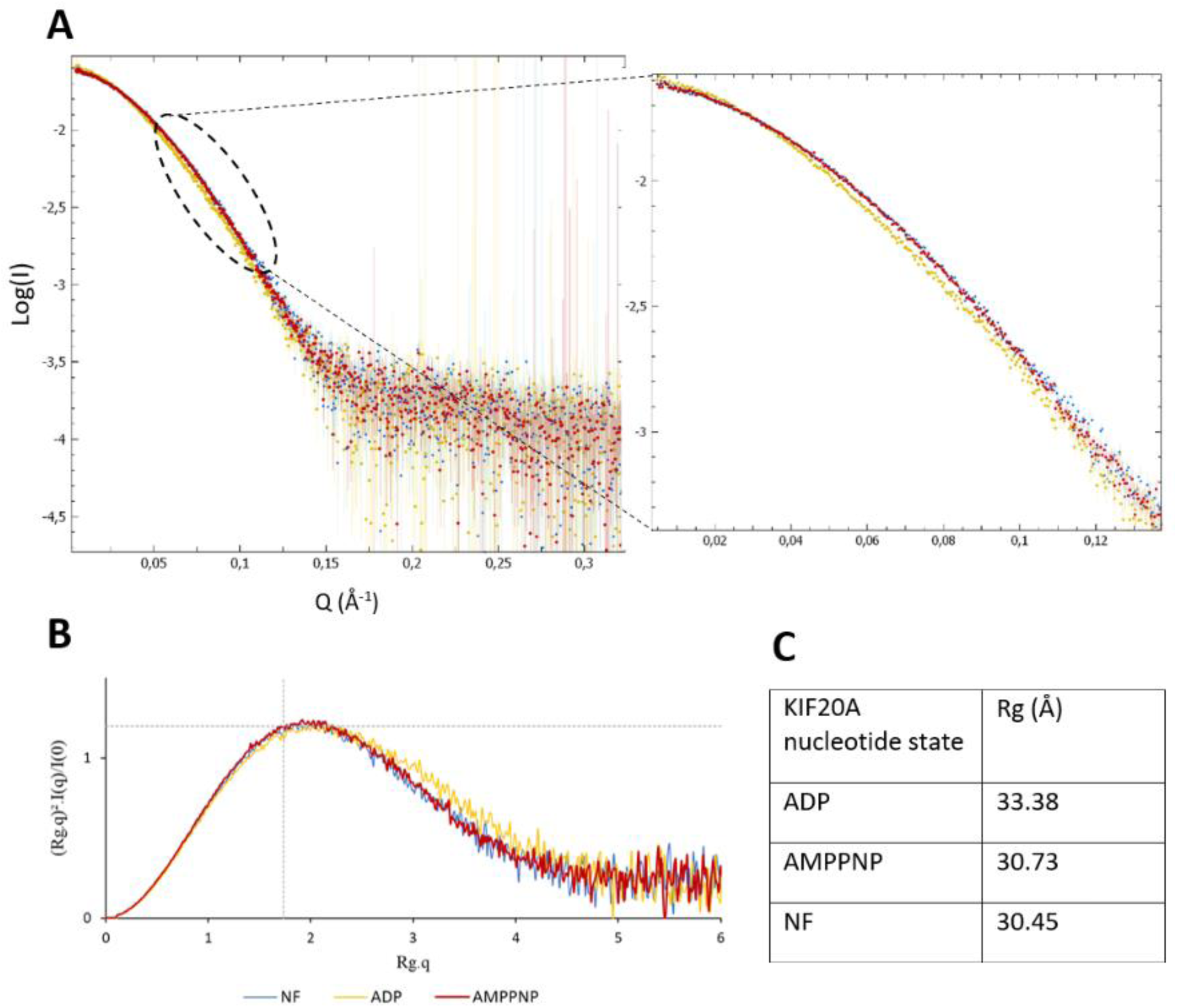
SAXS data of the 1-565 motor domain of KIF20A in different nucleotides state. showing that the ADP state shape is different than the others. In blue the NF curve, in yellow the ADP curve and in red the AMPPNP curve. **(*A*)** SAXS curve in logarithmic scale representation and zoom on the region at small Q showing the conformational differences in the overall shape of Kif20A. **(*B*)** Normalized Kratky representation of the different nucleotide states tested, **(*C*)** Rg values extracted from the Guinier region of the SAXS curves.

**Figure 4–figure supplement 2.**
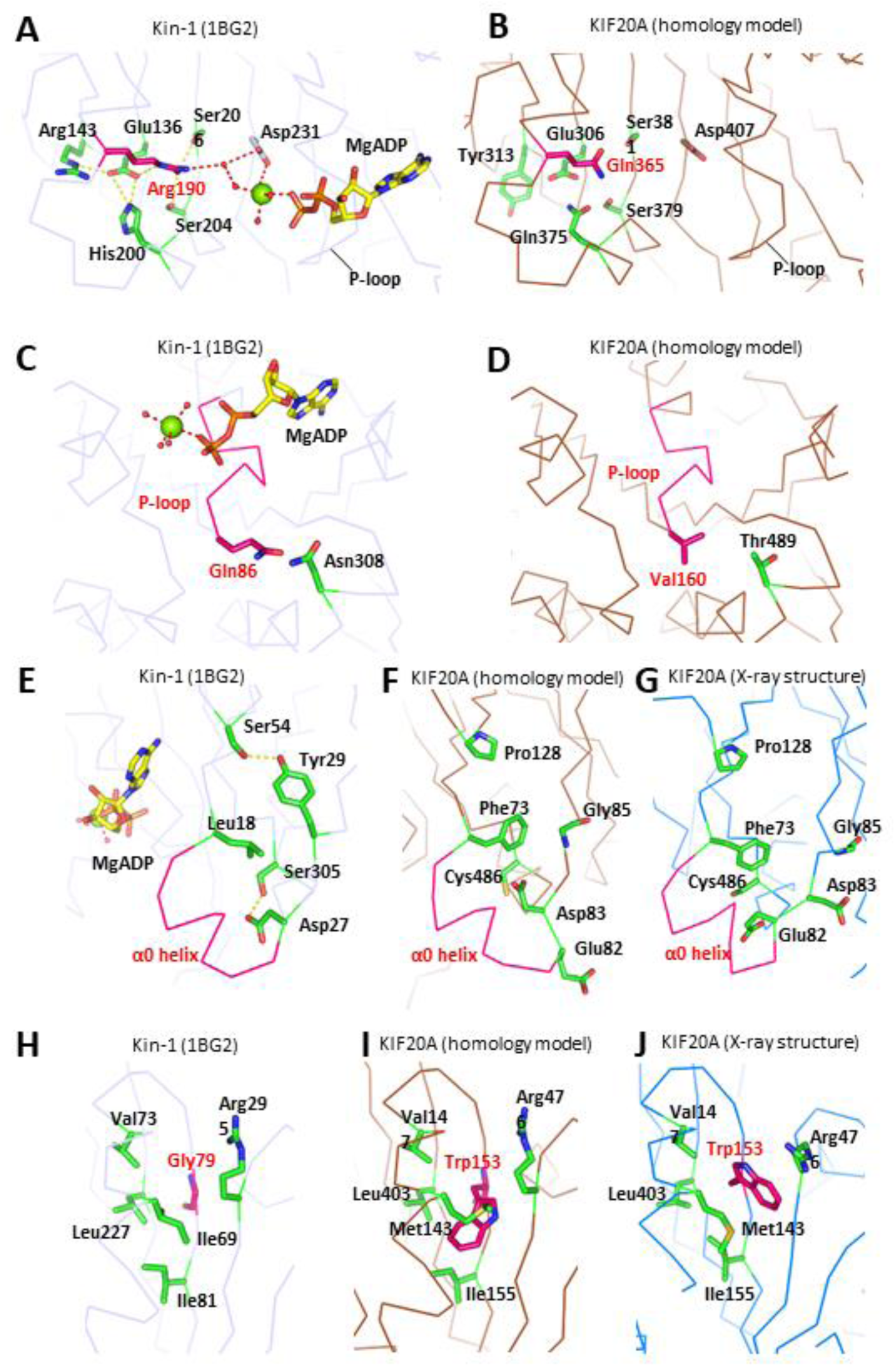
Some putative structural origins of the stability of the open NF state of KIF20A, revealed by homology modelling (KIF20A sequence was introduced in the 1BG2 Kin-1 structure). (*A, B*) Substitution of ^Kin-1^Arg 190 to ^KIF20A^Gln365. The interactions established by ^Kin-1^Arg 190 to stabilize the ADP-associated Mg^2+^ (A) cannot be formed in KIF20A, due to this Arg->Gln substitution, but also due to other substitutions of surrounding residues (^Kin-1^Arg143->^KIF20A^Tyr313 and ^Kin-1^His200->^KIF20A^Gln375) (B). (*C, D*) Substitution of the P-loop residue ^Kin-1^Gln86 to ^KIF20A^Val160. This substitution may lead to a less stable P-loop in KIF20A compared to Kin-1, thus may contribute to the observed lower affinity for MgADP in the former. (*E-G*) Outward positioning of the helix α0 in KIF20A. Some of the interactions that sustain the inward positioning of the α0 helix in Kin-1 (E) are different in KIF20A due to different residues at corresponding positions (homology model, F). The insertion of one residue (Glu82) in KIF20A sequence may also be involved in the particular positioning of this helix in KIF20A, since in the homology model the presence of this residue leads to a distortion of α0 helix (F). The position of α0 in 1BG2 is also characterized by an interaction between ^Kin-1^Ser54 and ^Kin-1^Tyr29 (E) that cannot be established by the equivalent ^KIF20A^Pro121 and ^KIF20A^Gly85 (F). Instead, the outward positioning of α0 in KIF20A is associated with a wider separation between these positions, which, therefore can contribute to the specific positioning of this helix (G). (*H-J*) Substitution of ^Kin-1^Gly79 to ^KIF20A^Trp153. In the homology model (I), the bulky ^KIF20A^Trp153 appears clearly incompatible with a twist of the central β-sheet similar to 1BG2, in which ^Kin-1^Gly79 occupies the equivalent position. Therefore, the presence of a tryptophan at this position appears more favorable to a high twist of the central β-sheet as observed in the open state of KIF20A (J).

**Figure 5–figure supplement 1.**
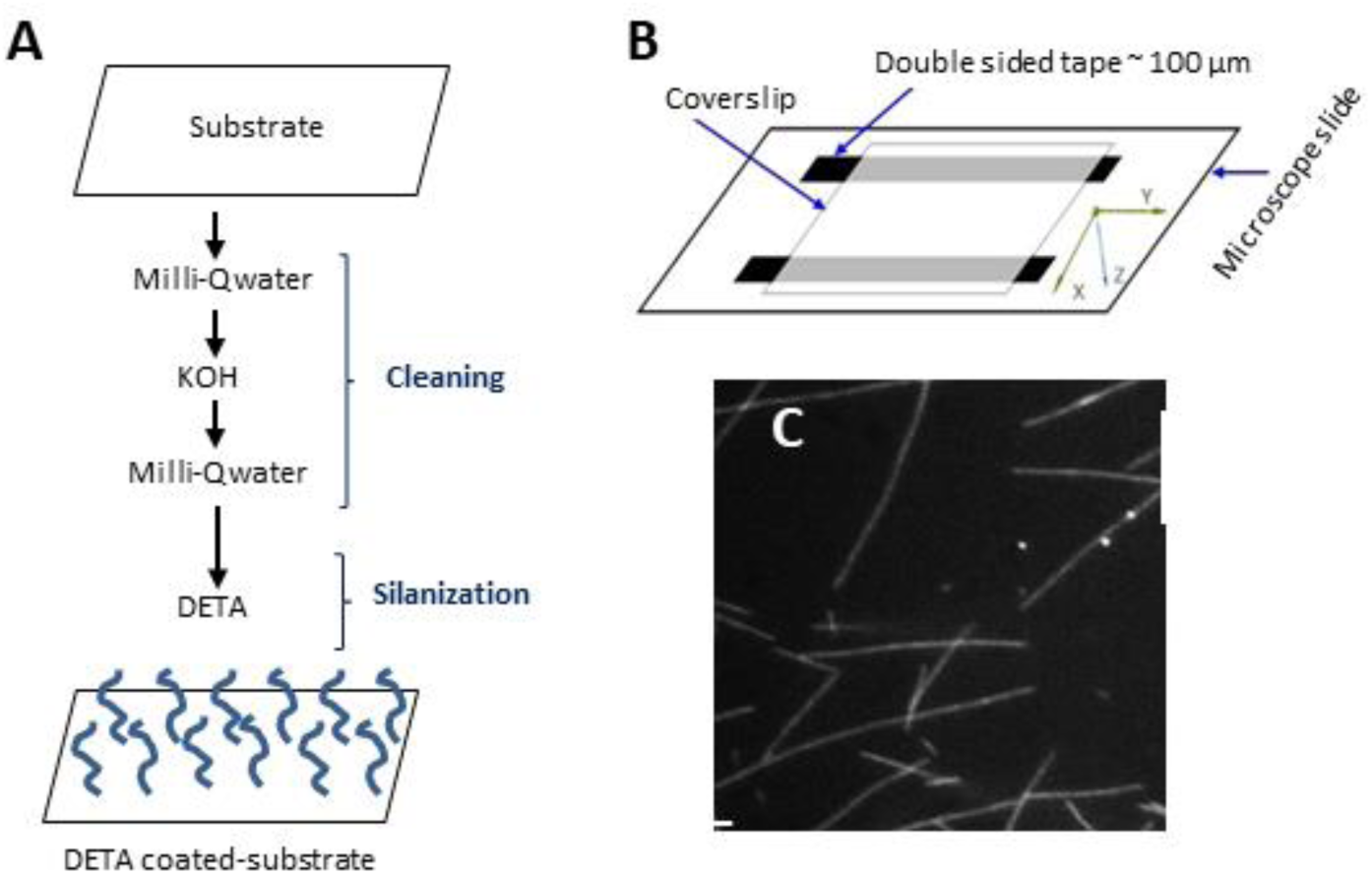
Sample chamber for motility assays, fluorescence intensity calibration. **(*A*)** Schematic of surface treatment: coverslips were cleaned with Milli-Q water and a KOH solution and then derivatized with a positively charged silane, 3-[2-(2-aminoethylamino)-ethylamino] propyl-trimethoxysilane (DETA). **(*B*)** Sample chambers were assembled using a microscope slide and a coverslip. Double-stick tape (thickness ~100 µm) was used as a spacer. Microtubules were flushed into the middle of the chamber, ~50 µm above substrate. **(*C*)** Tetramethyl-rhodamine-labeled microtubules on silanized coverslip (Scale bar: 10 µm).

**Figure 5–figure supplement 2.**
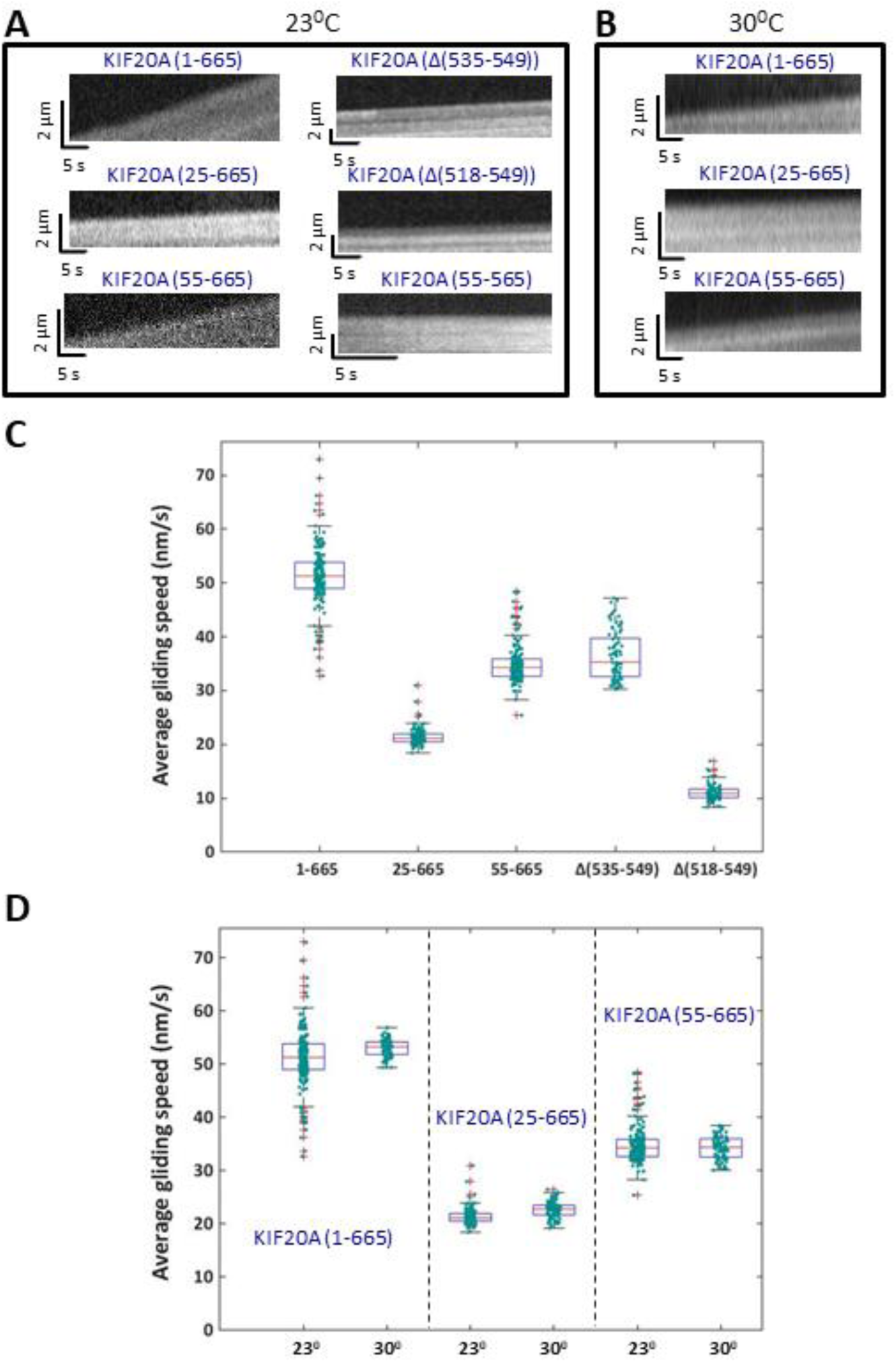
Gliding motility assays with various KIF20A constructs at 23°C and 30°C. **(*A*)** Kymographs of TMR-microtubules propelled by the KIF20A dimeric motor constructs 1-665, 25-665, 55-665, Δ(535-549), Δ(518-549) and the monomeric 55-565, respectively, at 23°C. The vertical axis of the kymographs represents microtubule displacement, the horizontal axis time. **(*B*)** Kymographs for selected constructs at 30°C. **(*C*)** Average gliding speeds produced by the respective constructs at 23°C. **(*D*)** Comparison of gliding speeds at 23°C and 30°C for selected constructs. Speeds increased by 2.7% for the 1-665 construct, by 5.3% for the 25-665 construct and by 0.8% for the 55-665 construct, which we consider insignificant in all cases. P-values derived from a two-tailed Student’s t-test for two samples with unequal variances to probe for statistical significance of the difference between the means of the distributions of speeds for a given construct at the two different temperatures were: 0.0655, 0.0004, 0.0493 for 1-665, 25-665 and 55-665, respectively, signifying that means at least two differences might be statistically relevant when assuming a threshold of P = 0.05, although more data would be needed to confirm these findings of potential small differences.

**Figure 5–figure supplement 3.**
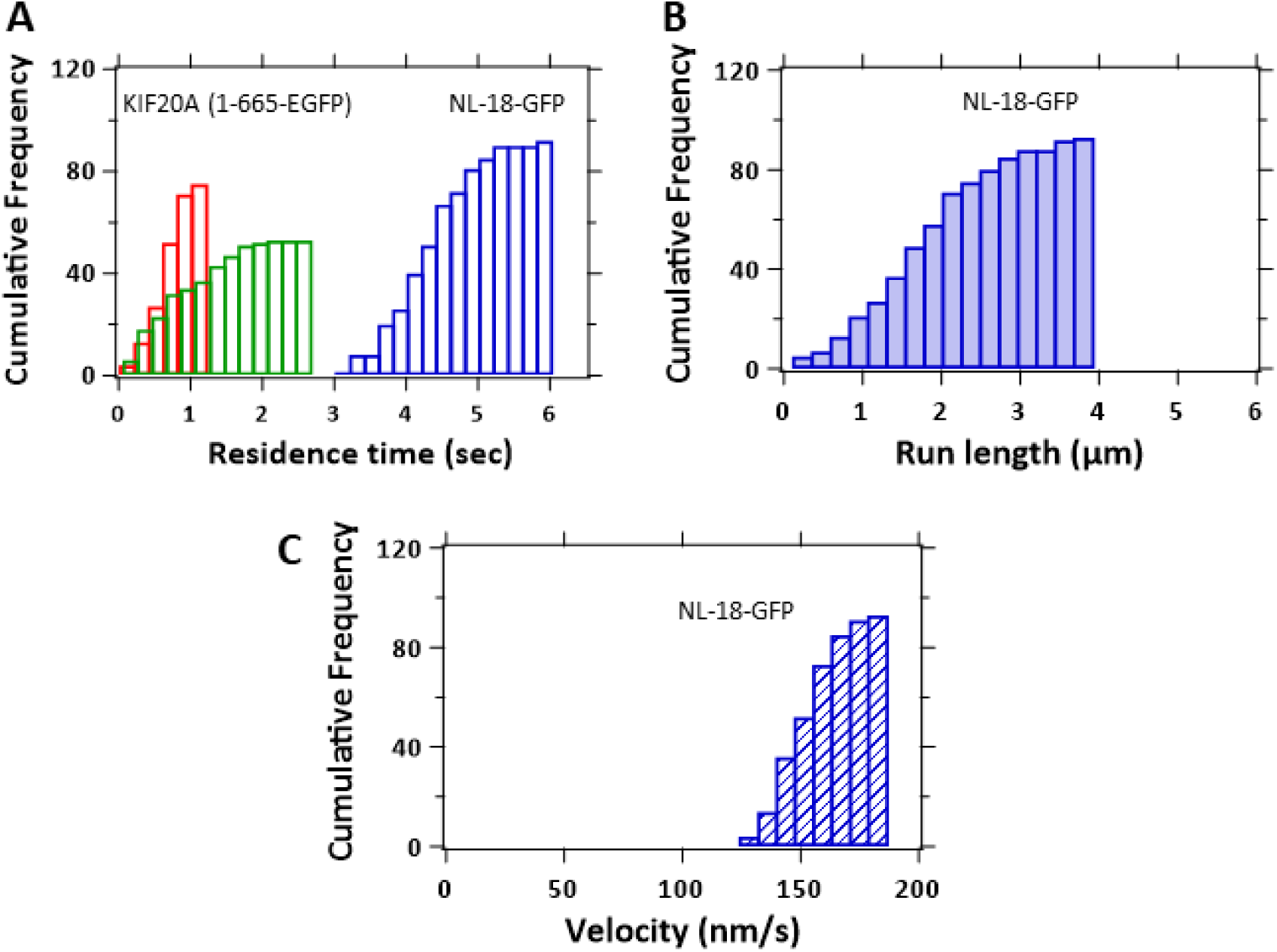
Single-molecule motility of 1-665-EGFP motor and kinesin-5-GFP (control). **(*A*)** Cumulative distribution functions of dwell times from single-molecule motility experiments with KIF20A 1-665 (red) and with Kin-5 (blue) as a control, measured from kymographs taken along microtubules (see Fig. 5 F and G). As control for purely diffusive dwells of KIF20A 1-665 in the detection volume, kymographs were evaluated from lines distant from any microtubule (green). Controls: Cumulative distribution functions for run lengths **(*B*)** and velocities **(*C*)**, measured from kymographs of GFP-labeled Kin-5 motors moving along microtubules (see Fig. 5F). Two independent experiments were performed with each motor protein.

**Figure 7–figure supplement 1.**
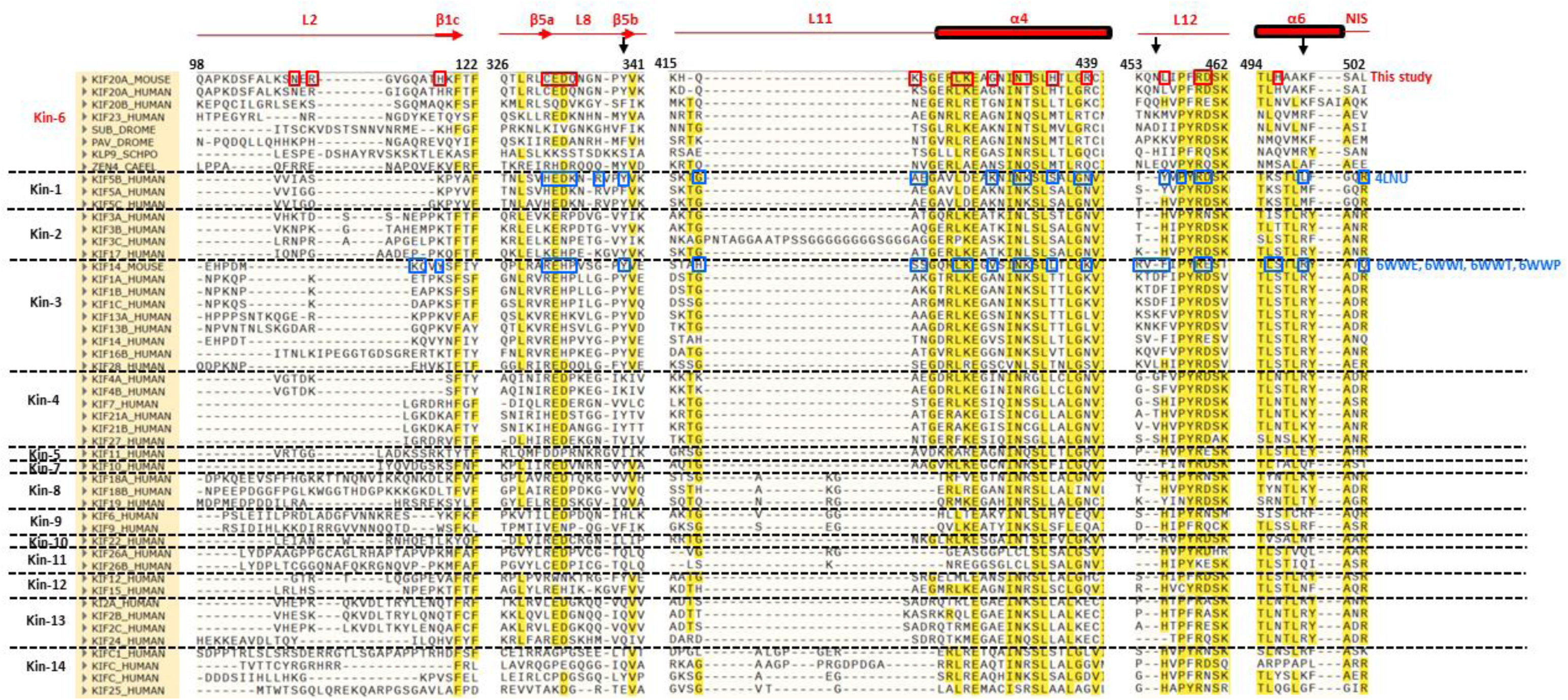
Alignment of areas of kinesin sequences that include KIF20A residues contacting microtubule. The residues within contact distance with α or β-tubulin from the either class 1 or class 2 structures of KIF20A-microtubule complex are boxed in red. By comparison, microtubule-contacting residues from the NF Kin-1-tubulin complex structure (4LNU) or NF KIF14-microtubule structures (6WWE, 6WWI, 6WWT or 6WWP) are boxed in blue. Several contacting residues are found in highly conserved areas are common in all available kinesin-microtubule or -tubulin complexes. Those residues are from L8, L11 and L12 loops, as well as from α4 and α6 helices. In addition, KIF20A and KIF14 both engage residues from their extended L2 loops in the interaction with microtubule, even though the sequence and the size of this loop are poorly conserved among kinesins. Arrows indicate residues not in contact with tubulin in KIF20A while their equivalent are part of the interface in Kin-1 or KIF14. In addition to the residues highlighted here, the class-1 KIF20A-microtubule structure reveals a contact between the KIF20A NL and microtubule through His513, Gly515 Ile516 and Pro517. The residues within contact distance between kinesin and tubulin were determined by a routine ‘findcontacts’ option in Chimera (Pettersen *et al*., 2004). The alignment was performed with the sequences of the kinesin structures discussed in this study and representative sequences for Kin-6s from different species and all kinesin classes from human, using the ‘Constraint-based Multiple Alignment Tool’ (COBALT) from the NCBI server (https://www.ncbi.nlm.nih.gov/tools/cobalt/re_cobalt.cgi).

**Figure 8–figure supplement 1.**
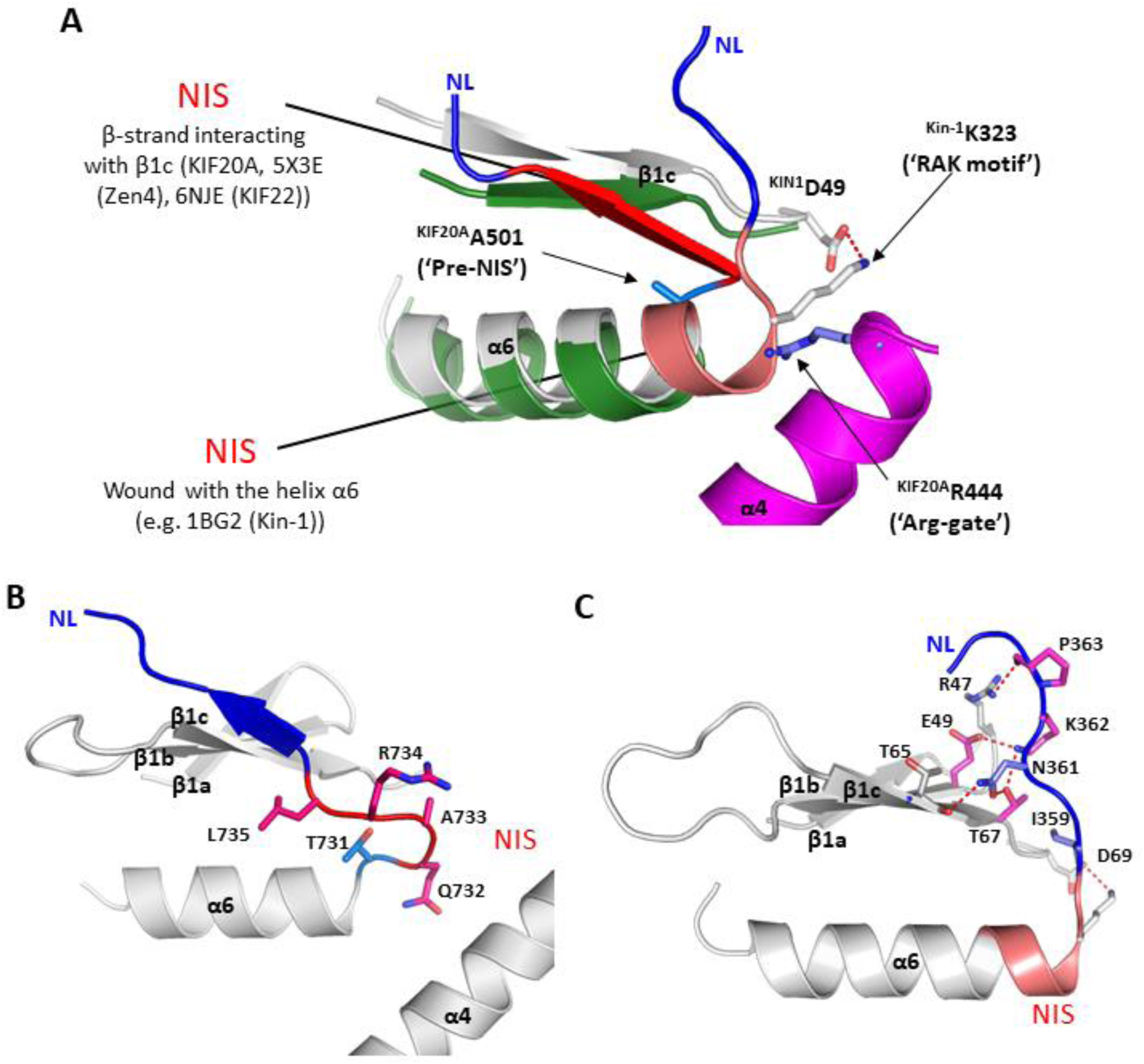
Involvement of the NIS in the orientation of the neck-linker and comparison with other plus-end kinesins. **(*A*)** Summary of the elements involved in the NIS backward docking in KIF20A, (the “pre-NIS residue Ala501 and the arginine gate residue Arg444, shown in blue) and of elements favoring the winding of the NIS with the α6 helix in Kin-1 (Lys323 residue from the “RAK motif” interacting with the conserved Asp49). **(*B*)** Backward orientation of the NL in the leading head of the 2-head-bound state of the Kin-3 KIF14 (6WWG) (Benoit *et al*., 2021). Constraints from the trailing head result in the stabilization of the proximal NL to interact with β1c similarly to that of the NIS in KIF20A. In this case, the NIS is also unwound, but it does not form an anti-parallel β-strand and rather forms a loop linking the α6 helix and the β-strand formed by the proximal NL. **(*C*)** Stabilization of the NL towards the microtubule minus end as observed in the ADP-bound Kin-5 structures (Goulet *et al*., 2014; Turner *et al*., 2001). In this conformation, the NIS is wound with the α6 helix, and the NL is stabilized on top of the β1a-β1c β-sheet involving multiple residues among which several residues are conserved in the Kin-5 class (magenta) but not in other classes (Turner *et al*., 2001).

**Figure 8–figure supplement 2:**
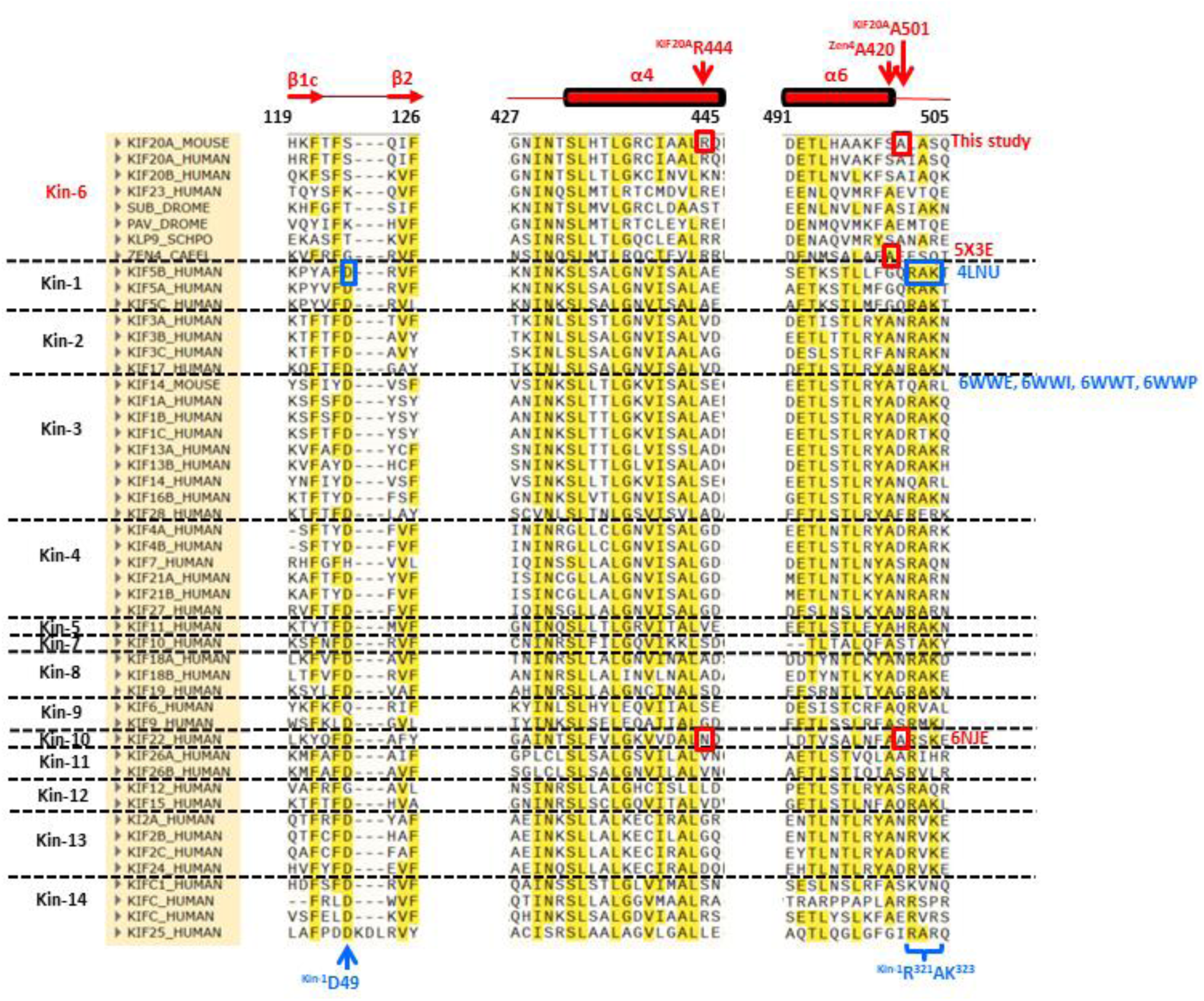
Alignment between several kinesin-6s with human kinesins of other classes showing conservation and variability of the elements guiding the backward docking. The Pre-NIS ^KIF20A^A501 that contributes to the backward docking in KIF20A via an incorporation into a specific hydrophobic pocket is shifted one residue earlier in Zen4, due to the insertion of a serine. In other kinesins, an alanine or Glycine residue is aligned with the Zen4 pre-NIS alanine that could play a similar role. However, in the KIF22 structure (6NJE) the residue playing this role is indeed equivalent to ^KIF20A^A501, suggesting that this position is more favorable to a backward docking of the NIS. The Arginine gate (^KIF20A^R444) is strongly conserved in Kin-6s but the equivalent position is rather occupied by small and hydrophobic residues in other kinesins, except in Kin-14s and in the Kin-10 KIF22 (6NJE), where the presence of a polar residue (Asn) participates in the unwinding of the NIS from the α6 helix and its backward docking, similarly to KIF20A. The ^KIF5B^R^321^AK^323^ motif in the NIS found at the end of the α6 helix in other pre-power stroke structures (with NIS wounded as an helix) and the structurally nearby residue ^KIF5B^D49 are mostly conserved in all kinesins except those of Kin-6, hence contributing to the NIS winding into α6 in these kinesins. In contrast, these resides are not conserved in Kin-6s, favoring the unwinding of the NIS.

**Figure 9–figure supplement 1:**
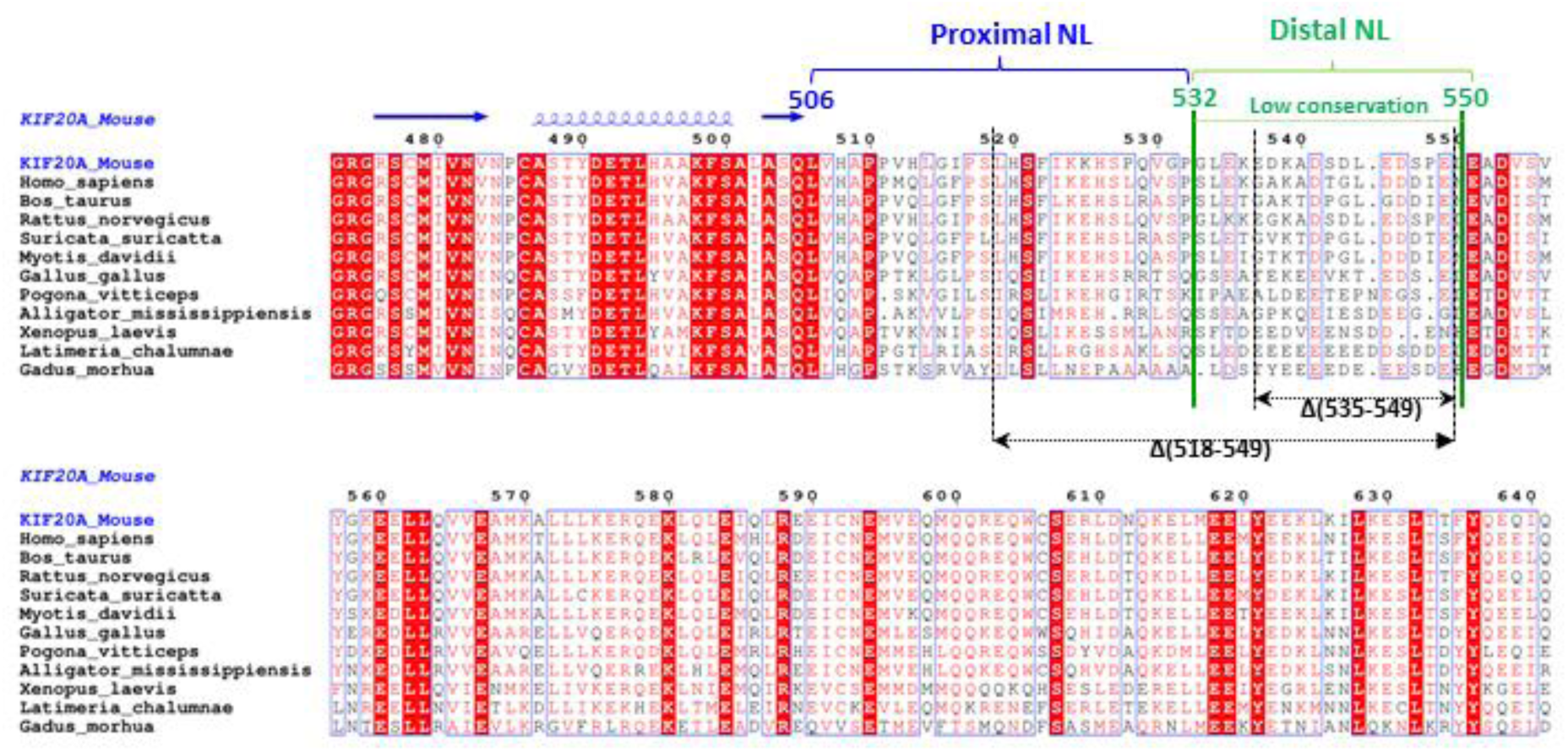
Alignment of the mouse KIF20A sequence used in this study with homologs from various species in the vicinity of the neck linker (506-553). The distal NL fragment 532-550 is less conserved among the species than the proximal NL fragment. Within this distal fragment, the deleted residues in the construct that exhibited unaffected gliding motility (Δ(535-549)) are indicated.

## Videos (separate files)

### Gliding motility assays

**Video 1.** Representative movie of a multi-motor gliding assay with 2 mM ATP in the assay buffer. TMR-labeled microtubules propelled by dimeric KIF20A 1-665 motors. Frame rate: 5 s^-1^, total recording time 80 s (sped up 20x).

**Video 2**. Representative movie of a multi-motor gliding assay with 2 mM ATP in the assay buffer. TMR-labeled microtubules propelled by dimeric KIF20A 55-665 (shortened N-terminus) motors. Frame rate: 5 s^-1^, total recording time 80 s (sped up 20x).

**Video 3.** Representative movie of a multi-motor gliding assay with 2 mM ATP in the assay buffer. TMR-labeled microtubules propelled by dimeric KIF20A 25-665 (shortened N-terminus) motors. Frame rate: 5 s^-1^, total recording time 80 s (sped up 20x).

**Video 4.** Representative movie of a multi-motor gliding assay with 2 mM ATP in the assay buffer. TMR-labeled microtubules propelled by dimeric KIF20A Δ(535-549) (shortened NL) motors. Frame rate: 10 s^-1^, total recording time 40 s (sped up 10x).

**Video 5.** Representative movie of a multi-motor gliding assay with 2 mM ATP in the assay buffer. TMR-labeled microtubules propelled by dimeric KIF20A Δ(518-549) (shortened NL) motors. Frame rate: 10 s^-1^, total recording time 40 s (sped up 10x).

**Video 6.** Representative movie of a multi-motor gliding assay with 2 mM ATP in the assay buffer. TMR-labeled microtubules propelled by dimeric Kinesin-1 (NK433) motors, control experiment. Frame rate: 10 s^-1^, total recording time 100 s (not sped up).

**Video 7.** Representative movie of a multi-motor gliding assay with 2 mM ATP in the assay buffer. TMR-labeled microtubules propelled by dimeric Kinesin-3 motors, control experiment. Frame rate: 5 s^-1^, total recording time 7.8 s (not sped up).

**Video 8.** Representative movie of a multi-motor gliding assay with 2 mM ATP in the assay buffer. TMR-labeled microtubules propelled by dimeric KIF20A 1-665-EGFP motors. Frame rate: 5 s^-1^, total recording time 80 s (sped up 20x).

### Single-molecule motility assays

**Video 9.** Single-molecule motility assay with dimeric KIF20A construct 1-665-EGFP. Frame rate: 10 s^-1^, total recording time 100 s (sped up 5x).

**Video 10.** Single-molecule motility assay with dimeric chimera Eg5kin-NL-18-GFP, control experiment. Frame rate: 10 s^-1^, total recording time 59.4 s (sped up 5x).

## Supplementary files

**Supplementary File 1.**
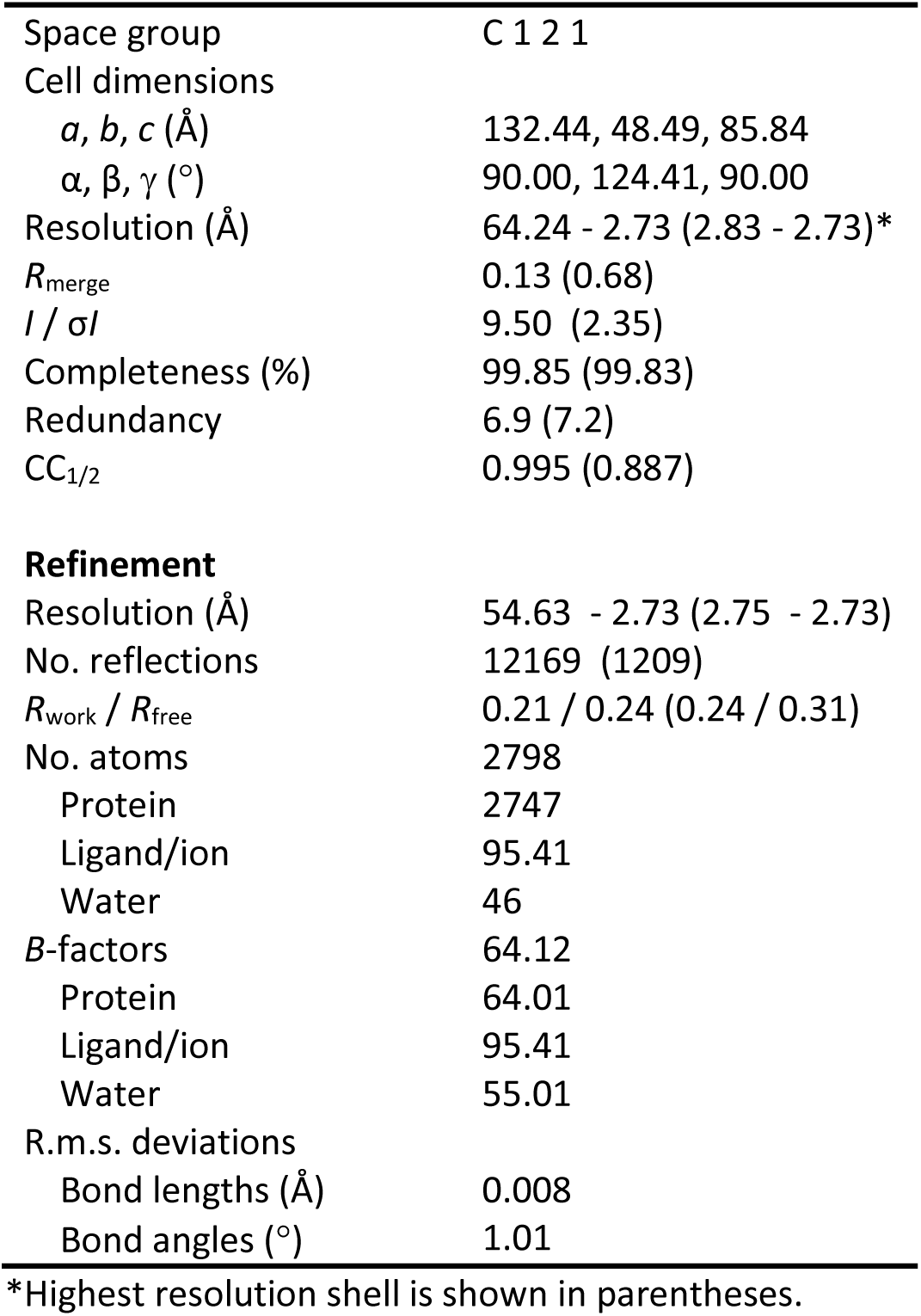
Statistics for X-ray data and model (PDB 8BJS)

**Supplementary File 2.** Cryo-EM data collection, refinement and validation statistics.

|  | Microtubule-KIF20A<br>nucleotide free<br>**Class-1**<br>(EMD-28789)<br>(PDB 8F1A) | Microtubule-KIF20A<br>nucleotide free<br>**Class-2**<br>(EMD-28787)<br>(PDB 8F18) |
| --- | --- | --- |
| **Data collection and processing** |  |  |
| Magnification (actual) | 105000 | 105000 |
| Voltage (kV) | 300 | 300 |
| Electron exposure (e<sup>-</sup>/Å<sup>2</sup>) | 56.1 | 56.1 |
| Defocus range (μm)<sup>a</sup> | 1.1 ; 2.7 | 1.1 ; 2.7 |
| Pixel size (Å) | 0.844 | 0.844 |
| Symmetry imposed<sup>b</sup> | Helical | Helical |
| Rise (Å) | 5.62 | 5.62 |
| Twist (deg) | 168.09 | 168.09 |
| Particle images identified as 15R<br>symmetry (no.) | 1.9 10<sup>3</sup> | 1.9 10<sup>3</sup> |
| Particle images in helical<br>reconstruction (no.) | 1.9 10<sup>3</sup> | 1.9 10<sup>3</sup> |
| Single particles (no.)<sup>c</sup> | 2.8 10<sup>6</sup> | 2.8 10<sup>6</sup> |
| Single particles used (no.)<sup>d</sup> | 243559 | 119232 |
| Overall resolution (Å) | 3.1 | 3.2 |
| FSC threshold | 0.143 | 0.143 |
| Kinesin resolution (Å) | 3.4 | 4.0<sup>e</sup> |
| Tubulin resolution (Å) | 3.0 | 3.2 |
| **Refinement** |  |  |
| Model composition |  |  |
| Non-hydrogen atoms | 10189 | 9891 |
| Protein residues | 1278 | 1242 |
| Ligands | 4 | 4 |
| R.m.s. deviations |  |  |
| Bond lengths (Å) | 0.0055 | 0.0058 |
| Bond angles (°) | 1.12 | 1.13 |
| Validation |  |  |
| MolProbity score | 1.58 | 1.38 |
| Clashscore | 4.62 | 3.53 |
| Poor rotamers (%) | 0.45 | 0.00 |
| Ramachandran plot |  |  |
| Favored (%) | 95.04 | 96.41 |
| Allowed (%) | 4.96 | 3.51 |
| Disallowed (%) | 0.00 | 0.08 |

<sup>a</sup> Range of the per-particle defocus estimates. The range corresponds to 90% of the particles used, 5% of the particle defocuses are below and 5% are above this range.
<sup>b</sup> Symmetry imposed during the symmetrical refinement done before the local refinements (rise and twists provided for helical symmetry only).
<sup>c</sup> Total number of particles after symmetry expansion.
<sup>d</sup> Number of asymmetric units, corresponding to a kinesin motor bound to a tubulin dimer, used after 3D classification.
<sup>e</sup> Note that the resolution of that kinesin in this class is very heterogeneous, with the highest resolution near the microtubule interface where some side chains are well resolved, while the rest of the kinesin is at much lower resolution.

**Supplementary File 3.** Unmodeled regions in the crystal and cryo-EM (class-1) nucleotide-free KIF20A structures.

|  | **Fragment absent in final model** | **Location in the MD** |
| --- | --- | --- |
| X-Ray<br>(MD alone, 55-510) | Met54-Ser61 | N-terminus |
|  | Asn109-Gln115 | L2 |
|  | Gln232-Pro285 | L6 |
|  | Ser320-Arg325 | L8 |
|  | Ser369-Arg378 | L9 (Switch I) |
|  | Glu412-Asn430 | L11 (Switch II) |
|  | Gln447-Asn454 | L12 |
|  | Ala509-Pro510-6xHisTag | C-terminus |
| Cryo-EM (Class-1)<br>MD-microtubule<br>complex, 1-565) | Met1-Glu58 | N-terminus |
|  | Glu234-Asp286 | L6 |
|  | 518-565-6xHis Tag | C-terminus |

## Notes

### Competing Interest Statement

The authors have declared no competing interest.

